# Spatio-temporal dynamics of M_1_ and M_2_ macrophages in a multiphase model of tumor growth

**DOI:** 10.1101/2025.02.06.636877

**Authors:** Ioannis Lampropoulos, Panayotis Kevrekidis, Christos Zois, Helen Byrne, Michail Kavousanakis

**Affiliations:** School of Chemical Engineering, National Technical University of Athens, Iroon Polytechneiou 9, Zografou, Athens, 15780, Greece; Department of Mathematics and Statistics, University of Massachusetts Amherst, 300 Massachusetts Ave, Amherst, 01003, Massachusetts, United States of America; Department of Radiotherapy and Oncology, Democritus University of Thrace, Dimokritou 7A, Komotini, 68100, Greece; Mathematical Institute, University of Oxford, Wellington Square, Oxford, OX1 2JD, England; Ludwig Institute for Cancer Research, University of Oxford, Wellington Square, Oxford, OX1 2JD, England

**Keywords:** multiphase model, finite element method, macrophages, chemotaxis, immunotherapy

## Abstract

This study investigates the complex dynamics of vascular tumors and their interplay with macrophages, key agents of the innate immune response. We model the tumor microenvironment as a multiphase fluid, with each cellular population treated as a distinct, non-mixing phase. The framework also incorporates diffusible species that are critical for processes such as nutrient transport, angiogenesis, chemotaxis, and macrophage activation. Numerical simulations of our model show how phenotypic and spatial heterogeneity in the macrophage population arises and how such heterogeneity impacts a tumor’s growth dynamics. Finally, we propose an immunotherapeutic strategy based on the experimental agent vactosertib which promotes an anti-tumor macrophage phenotype. Our simulations demonstrate an increased density of anti-tumor macrophages over the period of a few months, followed by a relapse period where the tumor regains its original dynamics.

## 1 Introduction

Understanding interactions between cancer cells and the immune system is a challenging research topic. Cancer can manipulate a host’s immune system to avoid eradication and even cooperate with it [1, 2]. In fact, cancer cells can remodel the immune system, causing cells to differentiate towards specific cell lineages, which promote tumor growth [1–3]. This ability contributes crucially to chronic inflammation, which, in turn, aids cancer growth and dissemination [2].

As highlighted in Dunn et al.’s seminal work [4], the immune system and cancer cells engage in continuous, dynamic interactions that shape tumor development. This complex process, termed *immunoediting*, is divided into three stages, often referred to as the three “E”s. The first stage, *Elimination*, occurs when the immune system successfully detects and destroys cancer cells. The second stage, *Equilibrium*, represents the phase in which the immune system manages to control tumor growth without completely eradicating it. Finally, in the *Escape* phase, some tumor cells adapt by developing mechanisms to evade immune detection and proliferate unchecked.

Among the first immune cells to mobilize after tumorigenesis are macrophages [1], products of monocyte differentiation [5–7]. Monocytes are found in the circulatory system and, as they extravasate into damaged tissues, they differentiate into macrophages [8]. During tumor growth, macrophages play pivotal yet contradicting roles. The significant variance in their functions stems from the dynamic nature of their phenotype. Macrophages can adopt a spectrum of phenotypes in response to their microenvironment [9–11]. Two prominent phenotypes are classically activated macrophages, M_1_, and alternatively activated macrophages, M_2_. The former exhibit pro-inflammatory characteristics and inhibit tumor progression, while the latter tend to be anti-inflammatory and promote tumor progression [10, 12]. Consequently, lower ratios of M_1_ to M_2_ are associated with poorer prognosis for cancer patients [13].

Numerous molecules are involved in shaping macrophage phenotype, and thereby, behavior. While considering all signaling molecules is beyond the scope of this study, it is important to examine the roles of certain key factors. Consider for example colony stimulating factor-1 (CSF-1), which is secreted by both cancer cells and tumor associated macrophages (TAMs). CSF-1 binds to CSF1R receptors which are highly expressed on the surface of macrophages and biases their movement towards the tumor. Effectively, CSF-1 can be viewed as a macrophage chemoattractant. At the same time, in response to environmental cues such as hypoxia, TAMs express cytokines such as the epidermal growth factor (EGF). Cancer cells which express EGF receptors, then migrate towards the TAMs, completing a paracrine loop, which facilitates cancer cell movement and may contribute to migration and metastasis [14–16].

TAMs contribute to tumor metastasis, as they attract cancer cells towards the vasculature via EGF-mediated paracrine signaling [14–17]. C-X-C motif chemokine 12 (CXCL12), also known as stromal cell-derived factor 1 (SDF-1), stimulates migration in most leukocytes [18], including TAMs. As a result, CXCL12 secreted by perivascular fibroblasts coupled with EGF-mediated paracrine signaling, drives migration of M_2_ macrophages and tumor cells towards the vasculature to promote metastasis [17, 17].

Macrophage behavior within a diseased tissue depends on its phenotype. Several factors determine a macrophage’s phenotype, including molecular signals and environmental conditions [19]. For example, transforming growth factor beta (TGF-*β*) drives macrophage polarization towards the M_2_ (pro-tumor) phenotype. As such, TGF-*β* represents a natural target for immunotherapy [17, 20].

Another important mediator is vascular endothelial growth factor (VEGF). VEGF is a diffusible cytokine, secreted by cancer cells under hypoxia. It stimulates mature endothelial cells to shed their protective pericyte layer and to proliferate. VEGF also maintains vascular networks in an immature state, where they lack proper structure and are prone to leaks and occlusion [21, 22]. Like tumor cells, TAMs secrete VEGF under hypoxia. This stimulates angiogenesis, providing the tumor with increased access to nutrients and facilitating metastasis [23].

Numerous computational studies have attempted to unravel the mechanisms underlying macrophage involvement in cancer growth. Initial modeling efforts focused on their anti-cancer pro-inflammatory behaviour [24–27], and their potential to serve as carriers for therapeutic agents [28–31]. More recent, models have focused on the phenotypic plasticity of macrophages and their ability to perform different functions depending on their microenvironment. Several ordinary differential equation (ODE) models have been developed to investigate macrophage polarisation [32–35]. The role of macrophages in diseased sites has also been studied using multi-scale models [36–38].

Regarding macrophage interactions within their micro-environment, Knútsdóttir et al. [39] studied various paracrine and autocrine loops, including the CSF-1/EGF loop discussed earlier, in order to understand how macrophages enable metastasis in breast cancer. Norton et al. [40] developed an agent-based model demonstrating that macrophages promote and facilitate cell migration and metastasis. Their findings suggested that invasive tumor cells rely on recruited macrophages to sustain their metastatic potential. Finally, Bull and Byrne [41] utilized an agent-based model to validate a novel weighted pair correlation function to investigate the spatio-temporal evolution of interactions between cancer cells and macrophages, study macrophage phenotype switching, paracrine/autocrine signaling and metastasis. In their study they showed how varying the rate of macrophage extravasation and macrophage chemotactic sensitivity to CSF-1 affect the ability (or inability) of macrophages to eliminate a tumor.

Tumor growth modeling that treats the tissue as a continuum has been the focus of numerous scientific studies, even when macrophages were not the primary focus. These works have significantly influenced the literature and shaped the direction of scientific research [42–46].

In this study, we present a multiphase model in a two-dimensional domain. Our model focuses on the complex behavior of macrophages within tumors. Naturally, the trade-off is that the present model does not provide the detailed local information that, e.g., the agent-based approach of [41] allows. It accounts for macrophage recruitment through the vasculature and the paracrine signaling that induces monocyte extravasation. Furthermore, the model tracks phenotype switching of macrophages from a pro-inflammatory to a pro-tumor state. In addition, the model investigates the role of cancer cells’ secretions and environmental factors -such as hypoxia- on alternative activation. We also investigate the impact of TAMs on tumor invasion by monitoring changes in the tumor’s growth rate in the presence of macrophages. Another feature of our model is the representation of hypoxic niches within the tumor and the simulation of TAM concentrations in these hypoxic regions.

The aim of immunotherapy is to enhance the immune system’s ability to recognize and eliminate cancer cells. Immunotherapeutic strategies include checkpoint inhibitors, CAR-T cell therapy, and cancer vaccines, which are designed to reinvigorate the immune system and re-establish immune surveillance against cancer [47]. We use our model to investigate the effect of immunotherapy that inhibits TGF-*β*/TGF-*β*R signaling on classically activated macrophages, motivated by the compound vactosertib. This drug binds to TGF-*β* receptors on macrophage surfaces, preventing the binding of TGF-*β* and, consequently, alternative activation. This mechanism effectively “freezes” the macrophage phenotype in a pro-inflammatory state [48, 49]. Vactosertib is currently undergoing clinical trials and has shown promise as an immunotherapeutic agent [50–53]. This therapy reprograms macrophages by blocking the downstream signaling pathways responsible for activating the pro-tumor phenotype [10, 54–57], rather than eliminating them or targeting other signaling pathways such as CSF-1/CSF-1R [54, 58–60]. In our study, we simulate vactosertib’s effect on macrophage behavior, and the subsequent effect on tumor growth. We monitor the immune response that is achieved with the help of vactosertib and compare it to the therapy-free scenario. Lastly, we consider the impact certain kinetic parameters have on the drug’s therapeutic efficacy.

Our paper is structured as follows: Section 2 presents the model developed for an untreated tumor scenario. Section 3 describes computational methods employed, including technical details about the numerical solution of the developed system of equations. Section 4 presents our findings, including the influence of macrophages on tumor growth, and the spatio-temporal distributions of TAMs within the tumor microenvironemnt. Section 5 introduces the model for a tumor treated with immunotherapy. Section 6 explores the impact of immunotherapy based on our findings. Finally, section 7 provides concluding remarks on the presented results and outlines potential directions for future research.

## 2 Therapy free model

We introduce a two-dimensional, multiphase model consisting of six interacting phases: healthy cells, cancerous cells and macrophages (M_1_ and M_2_), along with vascular and interstitial fluid phases. Each phase is modeled as a viscous fluid and characterized by the spatio-temporal evolution of its normalized concentration, denoted as *θ*_*i*_, with *i* representing each fluid phase:

- Healthy cells, *θ*_*h*_.
- Cancer cells, *θ*_*c*_.
- M_1_ macrophages, 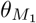.
- M_2_ macrophages, 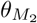.
- Vasculature, *θ*_*v*_.
- Interstitial fluid, *θ*_*int*_.

We associate with each phase a velocity vector, 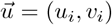, and a pressure term, *p*_*i*_, (*i* = *h, c, M*_1_, *M*_2_, *v, int*).

In addition to the fluid phases, the model accounts for several diffusible chemical species:

- Oxygen, *c*: a representative for nutrients in the system.
- VEGF, *g*: stimulates angiogenesis.
- CSF-1, *a*: mediates macrophage extravasation.
- CXCL12, *b*: acts as a chemoattractant for M_2_ macrophages.
- EGF, *l*: attracts cancer cells, facilitating tumor expansion and metastasis.
- TGF-*β, f*: polarises macrophages from M_1_ to M_2_.

We assume that the chemical species do not contribute to the system’s volume, as they are soluble. Oxygen is replenished via the vascular network, while the remaining diffusible species are expressed by specific cellular populations. Table 1 summarizes the variables defined in our system.

**Table 1.** List of the model’s variables.

| Variable name | Description | Variable name | Description |
| --- | --- | --- | --- |
| $\theta_h$ | Healthy cells concentration | $g$ | VEGF concentration |
| $\theta_c$ | Cancer cells concentration | $a$ | CSF-1 concentration |
| $\theta_v$ | Vessels concentration | $b$ | CXCL12 concentration |
| $\theta_{int}$ | Interstitial fluid concentration | $l$ | EGF concentration |
| $\theta_{M_1}$ | $M_1$ concentration | $f$ | TGF-β concentration |
| $\theta_{M_2}$ | $M_2$ concentration | $d$ | Drug concentration |
| $\theta_{M_1p}$ | Drug-bound $M_1$ concentration | $\vec{u}_i$ | Fluid phase, $i$, velocity |
| $c$ | Oxygen concentration | $p_i$ | Fluid phase, $i$, pressure |

A schematic of the model, depicting the interactions between the different cellular phases and chemical species is presented in Fig. 1.

**Fig. 1.**
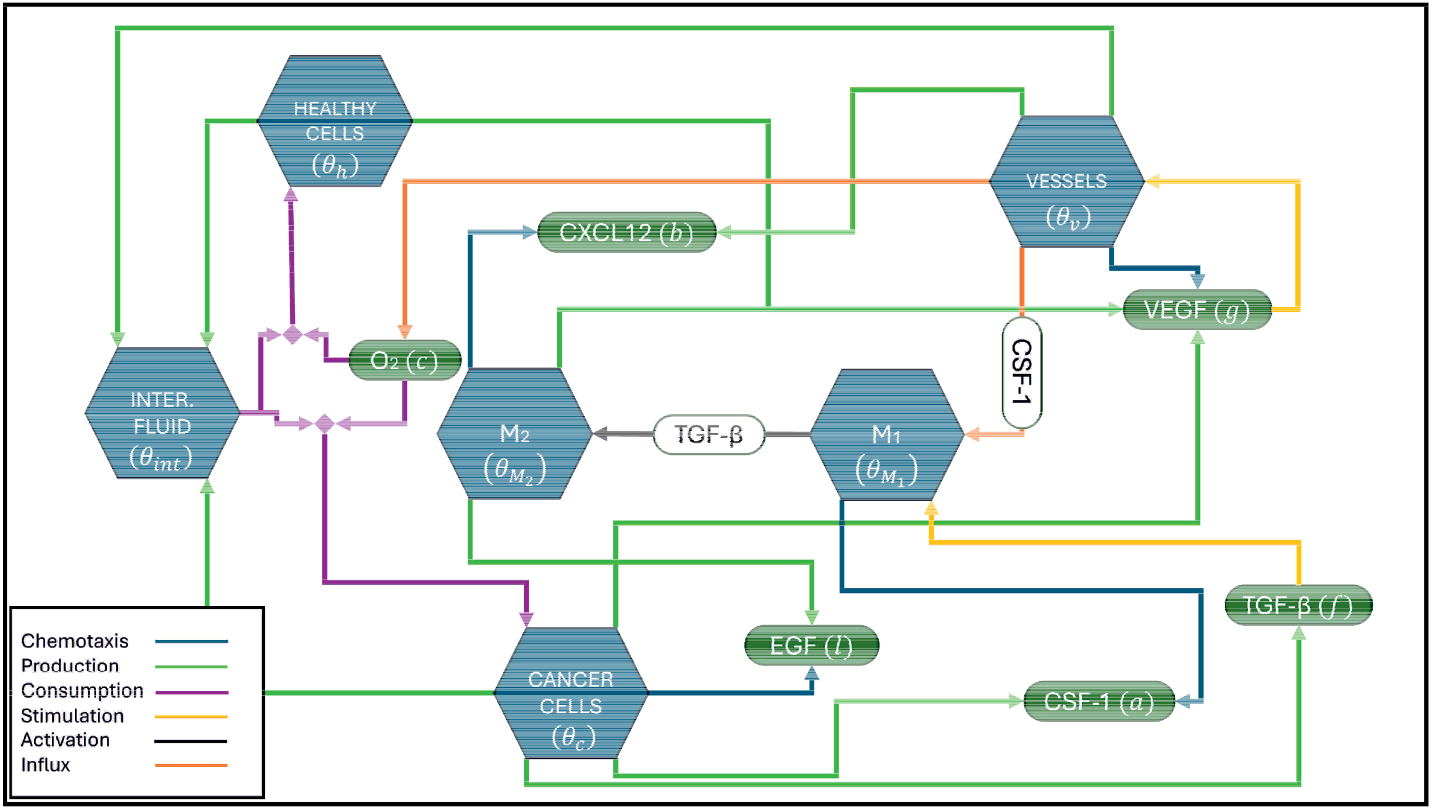
Model schematic. In the interactions presented, chemotaxis (and subsequent binding of molecules to the relevant receptors) is represented by blue arrows, production/secretion is illustrated by green arrows, consumption is shown as purple arrows, and influx through vasculature/extravasation is depicted by orange arrows.

### 2.1 Mass balance equations for cellular phases

It is commonly assumed that, at a macroscopic level, the density of living tissue is uniform [61]. As a result, the mass balance equations for the cellular phases are formulated in the following general form:

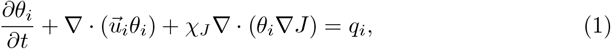

where *i* = *h, c, M*_1_, *M*_2_, *v, int* represents various phases, and *J* denotes the concentration of a chemical species, which acts as a chemoattractant for phase, *i*. In our model, there is at most one chemoattractant per cellular phase. Those pairings are listed in Table 2. The terms associated with mass transport appear on the left side of Eq. (1). In particular, 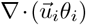 represents mass transport through convection, while *χ*_*J*_ ∇ · (*θ*_*i*_ ∇*J*) represents mass transfer of cellular phase, *i*, due to chemotaxis of spatial gradients of cytokine, *J*. The parameter *χ*_*J*_ denotes the strength of chemoattraction between cellular phase, *i*, and chemoattractant, *J*. Finally, the term *q*_*i*_ denotes the net production term for phase *i*. Not every chemokine attracts every cell type. Figure 2 illustrates the production and targets of our system’s chemoattractants, offering a summary of the above.

**Table 2.** Chemokine sensitive cells and their designated chemokine.

| Cell/Fluid phase | Chemokine/Chemical species | Source |
| --- | --- | --- |
| Cancer cells ($c$) | EGF ($l$) | [62] |
| Vasculature ($v$) | VEGF ($g$) | [21, 22] |
| Pro-inflammatory macrophages ($M_1$) | CSF-1 ($a$) | [63] |
| Pro-tumor macrophages ($M_2$) | CXCL12 ($b$) | [17, 18] |

**Fig. 2.**
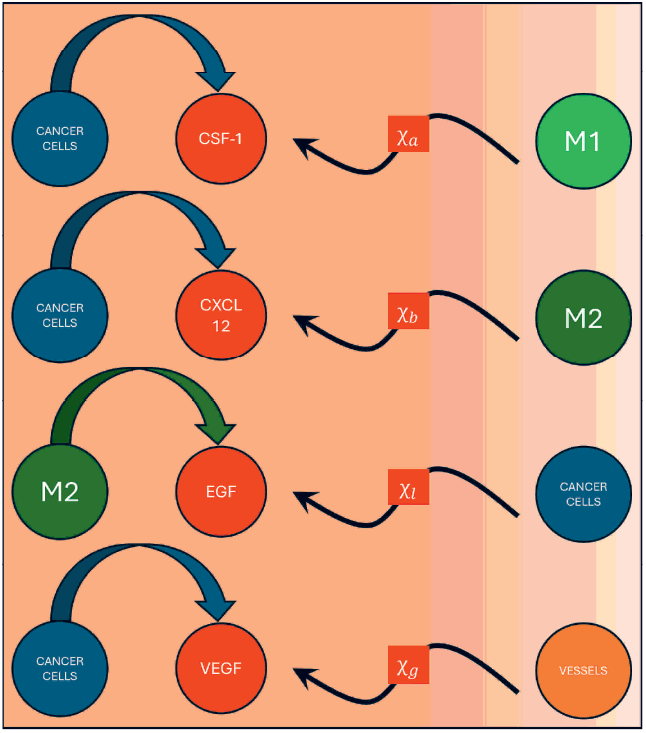
A summary of the chemokines included in our model, along with the cells that secrete them and the cells which are chemosensitive to them.

Boundary conditions for the mass balance equations (Eq. (1)) are specified on inflow segments of the tissue boundary (∂Ω). Inflow segments are defined as sections of ∂Ω where 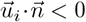, with 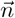 denoting the unit normal vector to the boundary. On inflow boundary segments, denoted ∂Ω^*inflow*^, we impose Dirichlet boundary conditions:

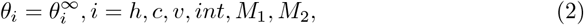

where 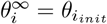, the concentration of species *i* at time *t* = 0. Homogeneous Dirichlet boundary conditions are imposed to simulate the state of a healthy tissue outside the domain of interest, Ω.

### 2.2 Production terms for mass balance equations

We now present the production terms, *q*_*i*_ (*i* = *h, c, v, int, M*_1_, *M*_2_), for the volume occupying phases.

#### 2.2.1 Healthy cells

We assume the production term for the healthy tissue surrounding the tumor can be written as:

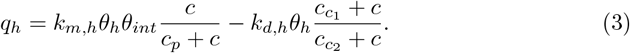

In Eq. (3), the first term simulates cellular proliferation and the second term represents cell death. We assume that healthy cells proliferate at a rate which is an increasing saturating function of the oxygen concentration, *c*, with a maximum value *k*_*m,h*_. The parameter *c*_*p*_ is the oxygen concentration at which the proliferation rate is half maximal. The proliferation rate is proportional to the interstitial fluid concentration, *θ*_*int*_ and the healthy cells concentration, *θ*_*h*_. The second term models cell death due to apoptosis and nutrient scarcity. We assume that it is a decreasing function of oxygen concentration, *c*, and, hence, that the parameters 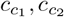 satisfy 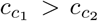. Further, *k*_*d,h*_ represents the basal rate of healthy cells death, attained in the limit *c* → +∞

#### 2.2.2 Cancer cells

The cancer cells proliferate and die, like the healthy cells. Additional terms are included to account for the boost in cancer cell proliferation caused by the presence of M_2_ macrophages and the increase in cancer cell death caused by M_1_ macrophages killing their targets:

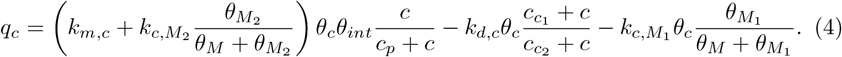

In Eq. (4), *k*_*m,c*_ denotes the maximum proliferation rate of cancer cells, and *k*_*d,c*_ is the rate at which cancer cells die. The cancer cell growth due to pro-tumor macrophage (M_2_) presence is included in the cancer cell proliferation term. 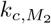 is the maximum increase in cancer cell proliferation due to the presense of M_2_ macrophages. *θ*_*M*_ is the concentration of macrophages where the rate of interaction between them and the cancer cells becomes half maximal. On the other hand, the killing of cancer cells caused by M_1_ macrophages is presented as a separate term. There, 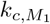 represents the maximum rate for the interactions between cancer cells and M_1_ macrophages, resulting to the death of the former. For simplicity, it is assumed that *θ*_*M*_ is common for interactions between cancer cells and both M_1_ and M_2_ macrophages. It is assumed that 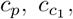, and 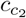 parameter values are identical for both healthy somatic and cancerous cells. Given the characteristics of malignant tumors consisting of rapidly proliferating and highly resilient cells, *k*_*m,c*_ *> k*_*m,h*_ and *k*_*d,c*_ *< k*_*d,h*_.

#### 2.2.3 Vasculature

The net production term for the vascular phase is given by:

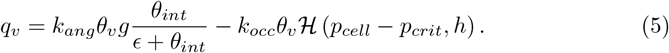

The first term in Eq. (5) describes angiogenesis, the process by which new blood vessels form from existing vessels [21]. We suppose that this rate is an increasing, saturating function of *θ*_*int*_ and proportional to the local concentration of VEGF, *g*. The interstitial fluid provides the material needed for endothelial cell proliferation. The parameter *k*_*ang*_, represents this phenomenon’s maximum rate, for a given VEGF concentration, *g*.

Tumor vessels are typically immature and prone to collapse [21]. We assume that blood vessels become occluded when the pressure exerted on them by the surrounding cellular phases exceeds the threshold value *p*_*crit*_. The value of *p*_*crit*_ represents the maximum pressure that the vessels can withstand. *p*_*cell*_ denotes the total pressure exerted by the cellular phases on the vessels:

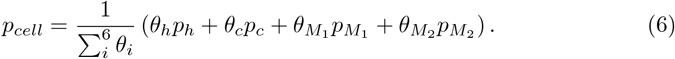

In Eq. (6), *p*_*h*_ and *p*_*c*_ denote the pressures exerted by the healthy and cancer cells and, similarly, 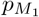 and 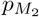 are the pressures exerted by the M_1_ and M_2_ macrophages. 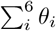 denotes the sum of concentrations of all cellular phases considered in the model, including the macrophage species. Finally, in Eq. (6), ℋ is a smooth approximation to the Heaviside function:

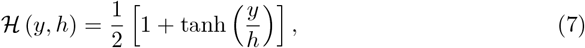

where the parameter *h* controls the smoothness (*h <<* 1).

#### 2.2.4 Macrophages M_1_ and M_2_

The production term for pro-inflammatory M_1_ macrophages is formulated as follows:

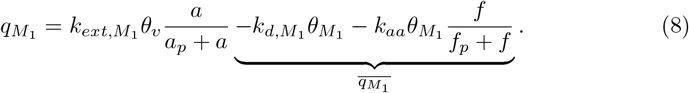

The first term represents the supply of macrophages from the vascular network. The macrophage extravasation rate is an increasing, saturating function of CSF-1’s concentration, *a*, with 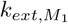 representing its maximum value. *a*_*p*_ is the CSF-1 concentration at which extravasation rate is half maximal. While there are reports of TAM proliferation in the tumor micro-environment, it is neglected in this study, motivated by experimental studies which show that most of the macrophages within the tumor microenvironment originate from differentiation of circulating monocytes [5, 54, 64].

We group the remaining two terms in 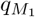 into a single sink term 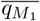 in order to distinguish them from the recruitment term which introduces mass to the system. The two sink terms represent: (i) macrophage death at a rate 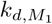, and (ii) alternative activation, the process by which M_1_ macrophages transition to a pro-tumor M_2_ phenotype. We assume that this process is an increasing, saturating function of TGF-*β, f*, with maximum rate *k*_*aa*_, and *f*_*p*_ being the TGF-*β* concentration at which the transition rate is half maximal.

The production term for M_2_ macrophages is similar to that of M_1_ macrophages:

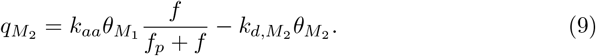

In Eq. (9), 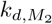 represents the death rate of M_2_ macrophages. We neglect source terms due to M_2_ proliferation and for M_2_ extravasation from vasculature.

#### 2.2.5 Interstitial fluid

The interstitial fluid serves as a passive medium that occupies the space between cells, known as the interstitium. It becomes enriched with necrotic material due to cell death, and absorbed during cell proliferation and angiogenesis. A fundamental modeling assumption is that the only external source of mass is due to extravasation of M_1_ macrophages. Unlike most multiphase models, the volume occupying phases do not sum to 1 (for *t >* 0) due to macrophage extravasation. Consequently, the production term for interstitial fluid, *q*_*int*_, is formulated as follows (balancing the rest of the contributions):

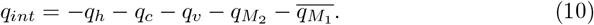

### 2.3 Momentum balance equations

We assume that the flow of the cellular species in the tissue has a Reynolds number *Re <* 1, and is characterized as creeping. Hence, inertial terms are neglected. The momentum balance for each cellular phase, *i*, is given by [44]:

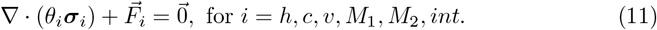

Here, ***σ***_*i*_ denotes the stress tensor of phase, *i*, and 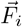 represents the forces exerted on phase *i* due to interactions with the other phases of the form. We view the volumetric phases as viscous fluids with stress tensors:

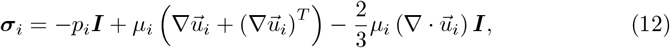

where *μ*_*i*_ denotes the dynamic viscosity of phase *i*.

The momentum source term 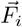 is formulated as follows:

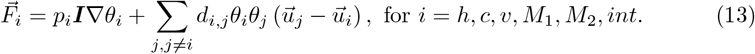

The first term captures the pressure’s, *p*_*i*_, effect on the phase’s surface, with ***I*** being the 2 *×* 2 identity matrix. Relative movement of phases *i* and *j* produces inter-phase drag as mentioned above, with drag coefficient *d*_*i,j*_.

We close Eq. (11) by imposing the following boundary conditions on ∂Ω:

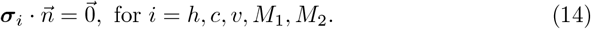

To ensure a unique solution, we also prescribe the velocity of the interstitial fluid on the boundary, ∂Ω, so that:

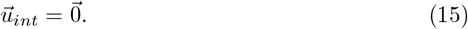

Having calculated the phase concentrations and velocities, it remains to determine the phase pressures. For this purpose, the continuity equation is formulated by summing the mass balance equation for each phase *i* (*i* = *h, c, v, M*_1_, *M*_2_, *int*), as formulated by Eq. (1):

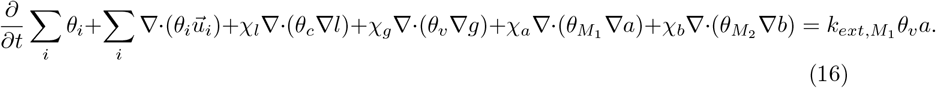

By computing *p*_*int*_ through Eq. (16), equations of state are formulated to relate the different pressures. Here, *p*_*v*_ is set equal to *p*_*ref*_ = 0 and for the remaining pressures, we define [44]:

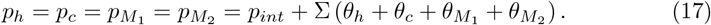

The function Σ(*θ*) is introduced to describe the pressure buildup due to local cell density exceeding the naturally established density in the tissue. If the natural cell density is denoted as *θ*^*^, function Σ is defined as follows:

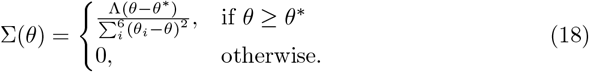

Here, Λ is a tension constant measuring the tendency of cells to restore their natural density.

### 2.4 Molecular species

Since the timescales for all processes involving molecular species (minutes) are short compared to those involving the cellular phases (days), all molecular species, *J*, are assumed to be in a quasi-steady state [42, 44, 65, 66]. The mass balance equations for the chemical species, *J*, reflect these considerations:

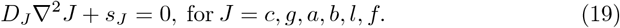

In Eq. (19), we assume that diffusion is the dominant mechanism for species transport and neglect transport due to advection. We denote by *s*_*J*_ the net source term for chemical species *J*. Diffusion is regulated by a diffusion coefficient *D*_*J*_.

The source term for oxygen, *s*_*c*_, is formulated as follows:

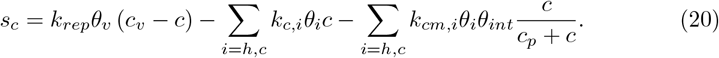

The first term represents the supply of oxygen from the vasculature. We assume that it is proportional to the concentration difference between *c*_*v*_ (the concentration of oxygen inside the vessels, which is assumed to be constant) and the oxygen concentration in the tissue, *c. k*_*rep*_ denotes the nutrient replenishment rate constant. The second term accounts for oxygen, *c*, consumption to sustain cells, where *k*_*c,i*_, (*i* = *h, c*) denotes the consumption rate for sustaining healthy cells (*θ*_*h*_) and cancer cells (*θ*_*c*_). The last term describes oxygen consumption for cell proliferation. It is assumed that it is an increasing, saturating function of *c*, with *k*_*cm,i*_, (*i* = *h, c*), denoting the maximum consumption rates for healthy and cancer cells, respectively. *c*_*p*_ is the oxygen concentration which results to the rate being half maximal.

For all the remaining chemical species, the source terms follow the same structure, consisting of a production term and two consumption terms: one corresponding to the molecule’s natural decay and one to the binding of the molecule its cognate receptors.

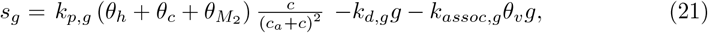

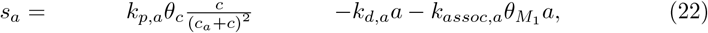

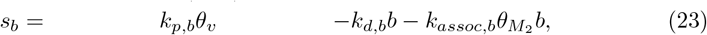

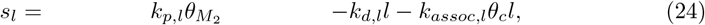

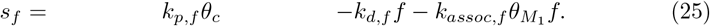

Here, *k*_*p,J*_, (*J* = *g, a, b, l, f*) denote the rates at which the chemicals are produced. *k*_*assoc,J*_ (*J* = *g, a, b, l, f*) represent the molecule consumption through binding on relevant receptors. *k*_*d,J*_ (*J* = *g, a, b, l, f*) denote the natural decay rates. VEGF, *g*, secretion is amplified as a response to hypoxia [21, 22], reflected on the quadratic term present in VEGF’s production term. It is assumed that healthy cells, cancer cells, and macrophages secrete VEGF at a rate which attains its maximum value 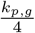 when *c* = *c*_*a*_ *< c*_*init*_ and decreases to zero as *c* → 0 and *c* → ∞. We assume that the rate at which cancer cells produce CSF-1 has the same biphasic dependence on the oxygen concentration. For simplicity, we assume that CXCL12’s, *b*, production is attributed to the vasculature since it is secreted by stromal cells surrounding the vasculature [17].

We impose Neumann boundary conditions for all molecular species, *J*, on the domain boundary ∂Ω:

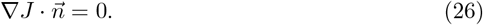

Before presenting numerical simulations, we non-dimensionalize our system. The characteristic time and length scales are chosen to be the doubling time for a healthy cell, which is roughly one day 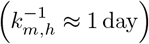 [67] and *L*_*o*_ = 400 *μm* respectively. The choice of characteristic length scale is based on the typical radii reached by cancer spheroids before angiogenesis commences [68–70].

Based on these two units we rescaled the system’s variables as follows:

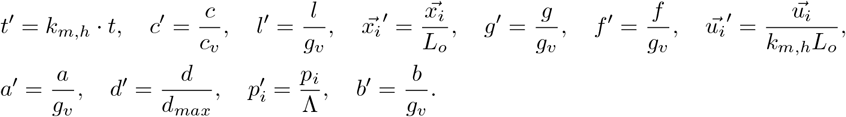

In addition to *k*_*m,h*_ and *L*_*o*_, several other units are used for the non-dimensionalisation of variables: *c*_*v*_ is the assumed-constant concentration of oxygen within the vessels; *g*_*v*_ is a typical concentration of VEGF, determined from the initial conditions described in Section 2.5. Λ is a tension constant measuring the tendency of cells to restore their natural density. *d*_*max*_ denotes the maximum concentration of the drug in the vasculature. Further details on the non-dimensionalisation can be found in SI, Chapter II.

The parameters utilized in the therapy-free model are summarized in Tables 3 and 4. Table 3 includes parameter values derived from related works and Table 4 contains those parameters that appear in the basic model and relate to the action of the M_1_ and M_2_ macrophages. All of the parameters are presented in dimensionless form, as described in Chapter II of the SI.

**Table 3.** Summary of model parameters, their descriptions, and estimated values [44, 66].

| Parameter name | Parameter value | Description |
| --- | --- | --- |
| $k_{m,c}$ | 2.0 | Cancer cell proliferation rate constant |
| $k_{d,h}$ | 0.15 | Healthy cell death rate constant |
| $k_{d,c}$ | 0.075 | Cancer cell death rate constant |
| $c_p$ | 0.25 | Proliferation rate saturation parameter |
| $c_{c1}$ | 0.2 | Death rate tuning parameter |
| $c_{c2}$ | 0.1 | Death rate tuning parameter |
| $c_a$ | 0.05 | Oxygen concentration yielding maximum VEGF and CSF-1 production rates |
| $k_{c,h}$ | 0.01 | Oxygen consumption rate constant by healthy cells for cell sustenance |
| $k_{c,c}$ | 0.01 | Oxygen consumption rate constant by cancer cells for cell sustenance |
| $k_{cm,h}$ | 0.1 | Oxygen consumption rate constant by healthy cells for mitosis |
| $k_{cm,c}$ | $k_{cm,h} \cdot k_{m,c}$ | Oxygen consumption rate constant by cancer cells for mitosis |
| $k_{occ}$ | 0.1 | Vascular occlusion rate constant |
| $p_{crit}$ | 0.3 | Critical pressure threshold |
| $\epsilon$ | 0.01 | Interstitial fluid's saturation parameter for angiogenesis |
| $h$ | 0.2 | Vascular occlusion smoothness parameter |
| $\mu_i$ | 10.0 | Dynamic viscosity for phase $i$ |
| $d_{i,j}$ | 1.0 | Drag between phases $i$ and $j$ |
| $\Lambda$ | 0.1 | cellular tension constant |
| $d_v$ | 1.0 | Oxygen diffusion coefficient |
| $D_g$ | 0.05 | VEGF diffusion coefficient |
| $k_{d,g}$ | 0.0264 | VEGF decay rate constant |
| $k_{assoc,g}$ | $6.57 \cdot 10^{-6}$ | VEGF association rate constant |

**Table 4.** List of parameters associated with the default model.

| Parameter name | Parameter value | Description | Source |
| --- | --- | --- | --- |
| $\theta_M$ | 0.1 | Cancer cell-macrophage interaction saturation constant | - |
| $k_{c,M_1}$ | 0.2 | Cancer cell - $M_1$ interaction rate constant | - |
| $k_{c,M_2}$ | 5.0 | Cancer cell - $M_2$ interaction rate constant | - |
| $k_{d,M_1}$ | 0.075 | $M_1$ death rate constant | [71] |
| $k_{d,M_2}$ | 0.075 | $M_2$ death rate constant | [71] |
| $k_{aa}$ | 0.5 | Alternative activation rate constant | - |
| $k_{ext,M_1}$ | 2.0 | Macrophage migration rate constant | - |
| $D_a$ | 0.11 | CSF-1 diffusion coefficient | [41] |
| $D_b$ | 0.001 | CXCL12 diffusion coefficient | [41] |
| $D_l$ | 0.11 | EGF diffusion coefficient | [41] |
| $D_f$ | 0.5 | TGF-β diffusion coefficient | [41] |
| $\chi_g$ | 0.14 | Chemotaxis constant for chemoattraction due to VEGF | SI Section I |
| $\chi_a$ | 0.14 | Chemotaxis constant for chemoattraction due to CSF-1 | - |
| $\chi_l$ | 0.14 | Chemotaxis constant for chemoattraction due to EGF | - |
| $\chi_b$ | $5 \cdot \chi_a$ | Chemotaxis constant for chemoattraction due to CXCL12 | - |
| $a_p$ | 0.25 | Macrophage migration saturation parameter | - |
| $f_p$ | 0.015 | Alternative activation saturation parameter | - |
| $k_{p,g}$ | 0.00959 | VEGF production rate constant | SI Section I |
| $k_{p,a}$ | 0.00959 | CSF-1 production rate constant | - |
| $k_{d,a}$ | 0.0216 | CSF-1 decay rate constant | [41] |
| $k_{assoc,a}$ | $8.75 \cdot 10^{-6}$ | CSF-1 association rate constant | - |
| $k_{p,b}$ | 0.54806 | CXCL12 production rate constant | SI Section I |
| $k_{d,b}$ | $0.2273 \cdot 10^{-2}$ | CXCL12 decay rate constant | [41] |
| $k_{assoc,b}$ | $7 \cdot 10^{-6}$ | CXCL12 association rate constant | - |
| $k_{p,l}$ | 0.00959 | EGF production rate constant | - |
| $k_{d,l}$ | 0.0216 | EGF decay rate constant | [41] |
| $k_{assoc,l}$ | $6.93 \cdot 10^{-6}$ | EGF association rate constant | - |
| $k_{p,f}$ | 0.00959 | TGF-β production rate constant | - |
| $k_{d,f}$ | 0.436 | TGF-β decay rate constant | [41] |
| $k_{assoc,f}$ | $7 \cdot 10^{-6}$ | TGF-β association rate constant | - |

### 2.5 Initial conditions

To establish the system’s initial conditions, we simulate the steady state condition of a healthy tissue. Consequently, we define the following conditions reflecting a healthy tissue’s homeostatic state:

- The volume fraction of healthy cells is set equal to 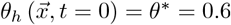.
- The tissue devoid of cancer cells: 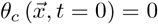.
- All present chemical species (oxygen and VEGF) are uniformly mixed.
- CSF-1, CXCL12, EGF, and TGF-*β* are molecules associated with cancerous inflammation and thus are absent in the healthy tissue.
- There is no inflammation and no inter-cellular stresses: 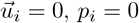, *p*_*i*_ = 0.
- Undifferentiated monocytes (macrophage progenitors) are present in extremely low concentrations within the tissue: 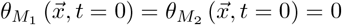.

As more thoroughly explained in SI Section I, the system can be considered closed. Consequently, the homeostatic tissue maintains a constant volume and the non-dimensionalised concentrations of the cellular phases are equivalent to their volume fractions. Based on this premise, we can simplify the calculation of *θ*_*int*_:

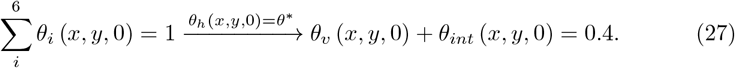

Combining the aforementioned considerations results in the following (non-dimensionalised) system:

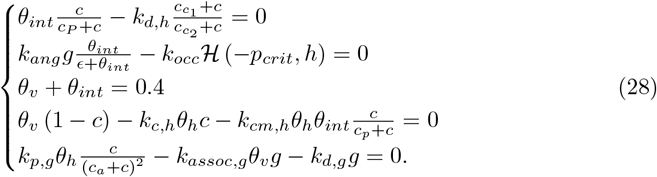

Solving the above yields the system’s equilibrium state. To compute a unique solution for the developed system, we consider *k*_*ang*_ as an unknown parameter. *k*_*ang*_ is a parameter whose value is challenging to determine experimentally so, it is considered an unknown parameter and solved for in Eq. (28): *k*_*ang*_ = 4.87 · 10^−3^. Initiating a cancerous seed centered at (*x, y*) = (0, 0) with a radius *r*_*o*_ = 1, results in the initial conditions for all utilized variables. The non-zero initial conditions are summarized in Table 5:

**Table 5.** Initial conditions of the model’s variables.

| Variable | Value/expression | Description |
| --- | --- | --- |
| $\theta_h$ | $0.6 - \theta_c(x, y, 0)$ | Healthy cells dimensionless concentration |
| $\theta_c$ | $\begin{cases} 0.05 \cos^2\left(\frac{\pi r}{2}\right), & \text{if } \sqrt{x^2 + y^2} \leq R_o = 1 \\ 0, & \text{otherwise.} \end{cases}$ | Cancer cells dimensionless concentration |
| $\theta_v$ | 0.0175 | Vessels dimensionless concentration |
| $\theta_{int}$ | 0.3825 | Interstitial fluid dimensionless concentration |
| $c$ | 0.25 | Oxygen's dimensionless concentration |
| $g$ | 0.6 | VEGF's dimensionless concentration |

## 3 Methods

Numerical simulations were performed using the Comsol Multiphysics ^®^software employing the Finite Elements Method (FEM). The computational domain is a circular disk, discretized with an unstructured mesh generated using Delaunay triangulation. The disk has a radius of 30 dimensionless units, and all computations are executed in non-dimensionalised units. Details of the non-dimensionalization process can be found in SI Section II.

All simulations are initiated by embedding one or more cancerous lesions within a healthy tissue. For a default simulation involving a tumor originating from single seed and presenting by both macrophage phenotypes in its micro-environment, the model solves approximately 250, 000 degrees of freedom. It takes approximately 100 hours of computational time on an AMD Ryzen 9 3900X 12-Core Processor to simulate 300 dimensionless time units (corresponding to approximately 300 days tumor growth).

## 4 Therapy-free model results

In this section, we present numerical results for the therapy-free model. We focus initially on the basic model which describes the growth of a small cluster of cancer cells located at the domain’s center. We then investigate the impact of varying key model parameters on the system dynamics, and finally present simulation results in which multiple tumor lesions merge. Since all computations are performed in dimensionless units, unless stated otherwise, all results are presented in dimensionless form without the need for the “′” and “*” symbols as presented in the SI.

A key feature of the model is its ability to determine the spatial distributions of each cellular phase and chemical species. In the left panel of Fig. 3, we present the distributions of cancer cells and macrophages at dimensionless times *t* = 150 and *t* = 250, for a tumor originating from a single cancerous lesion. Of particular interest is the ability of each macrophage phenotype to infiltrate the tumor. As expected, the distribution of M_1_ macrophages is broader than the distribution of M_2_ macrophages. The M_1_ macrophages concentrated around the tumor’s periphery. When they infiltrate the tumor, they are exposed to TGF-*β* which triggers alternative activation. Consequently, the M_2_ macrophages are concentrated beyond the tumor’s outer boundary.

**Fig. 3.**
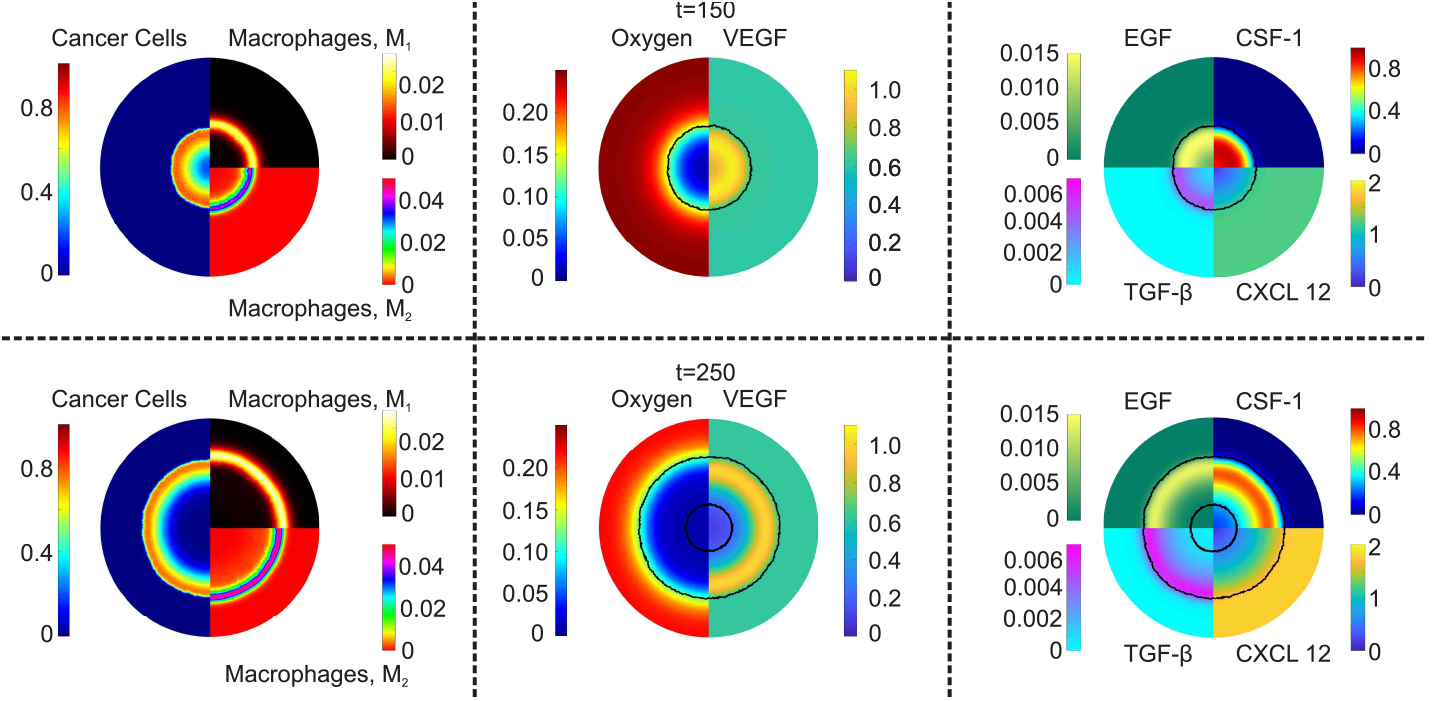
Basic model spatial distributions at dimensionless times: *t* = 150 (first horizontal row) and *t* = 250 (second row). The left panels depict the spatial distributions of the cancer cells and macrophages species: the left half-disk shows cancer cells, the right-upper disk depicts *M*_1_ macrophages and the right-bottom disk shows *M*_2_ macrophages. The middle panel shows the oxygen distribution (left half-disk) and the VEGF concentration distribution (right half-disk). The right panel shows the spatial distributions of EGF (top left quadrant), CSF-1 (top right quadrant), TGF-*β* (bottom left quadrant) and CXCL 12 (bottom right quadrant). The black lines denote the tumor’s boundaries.

Complementing the cancer and macrophage distributions, the middle and right panels of Fig. 3 depict the distributions of the chemical species, at *t* = 150 and *t* = 250. To provide context on their relative positions within the domain, we indicate the tumor’s boundary (and the necrotic boundary) as a black line. These boundaries are defined as the contour where *θ*_*c*_ = 0.01.

The plots of oxygen distribution (left half-disk in the middle panels of Fig. 3 show how the tumor’s expansion leads to vascular occlusion, which reduces tissue oxygenation. As the tumor grows, oxygen diffusion becomes insufficient to meet the tissue’s demands, and hypoxic areas form (light and medium dark blue). Hypoxia drives the production of VEGF and CSF-1, their concentrations peaking in the hypoxic regions. In contrast, concentrations of EGF and TGF-*β* are greatest within the tumor’s well-oxygenated, proliferative zone. The elevated cancer cell concentrations in this region enhance production of CXCL12. Due to its slow rate of diffusion, CXCL12 remains localized near its site of production.

Motivated by the numerical results for the basic model, we now investigate how macrophages can inhibit or facilitate tumor growth. We consider three scenarios: (i) a tumor where both macrophage phenotypes are present; (ii) a tumor where only the M_1_ phenotype is expressed, and (iii) a tumor where both M_1_ and M_2_ macrophages are present (they occupy space) but they are inert.

We compare the spatial distributions of cancer and immune cells for each case. In Fig. 4, we present the average radial distributions of cancer cells and macrophages at dimensionless times *t* = 100, 200, and 300. We observe that when both M_1_ and M_2_ macrophages are active, the cancer cells invade most rapidly. Secondly, the radial distribution of the M_1_ macrophages is broader as they enter the system via the vascular network and migrate towards the tumor. Conversely, M_2_ macrophages penetrate deeper into the tumor and localise in hypoxic regions, where they promote tumor cell proliferation.

**Fig. 4.**
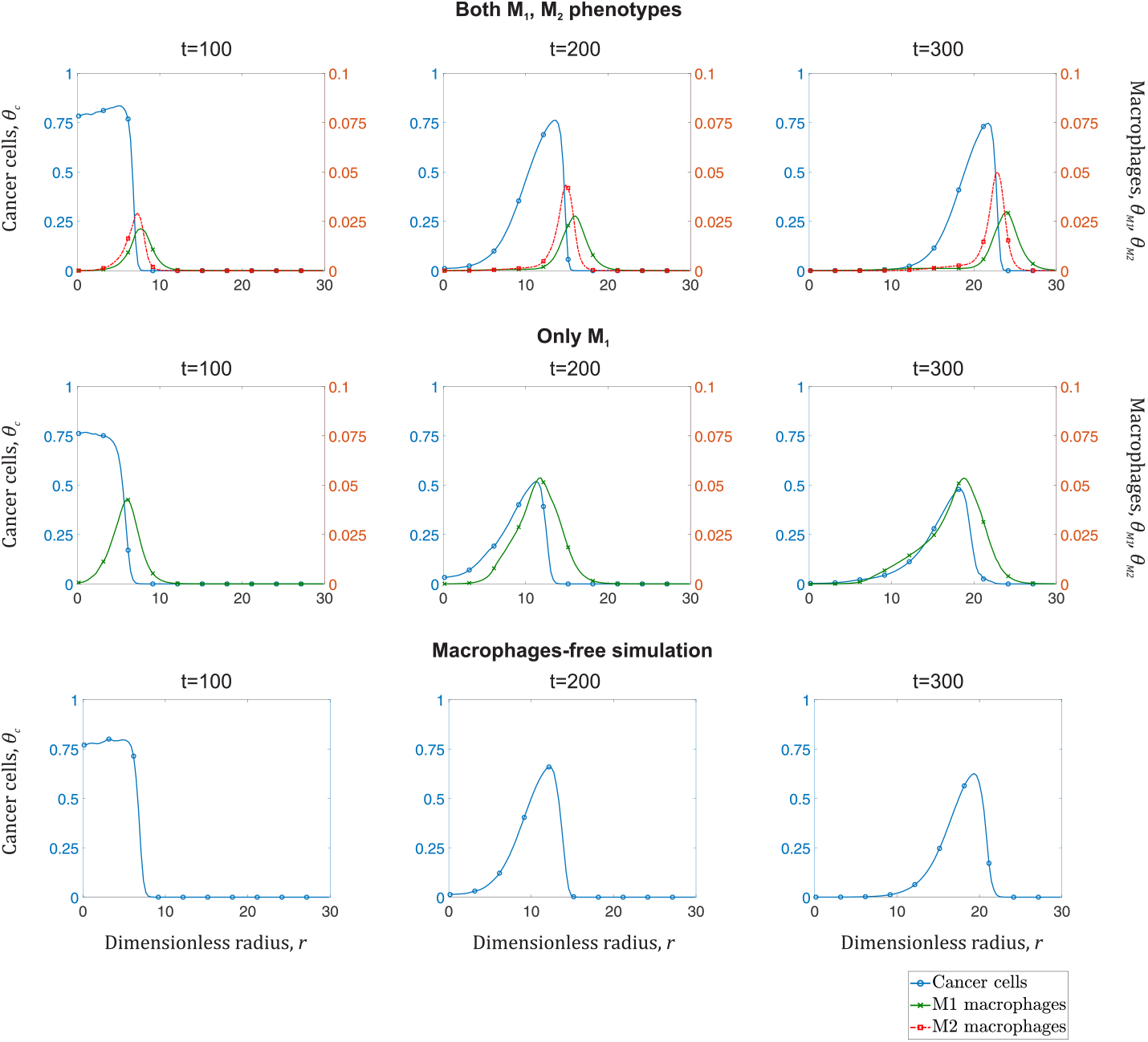
Series of plots showing the average radial distributions of cancer cells (left y-axis - blue curve with circles), M_1_ macrophages (right y-axis - green crossed curve), and M_2_ macrophages (right y- axis - red dotted curve with squares) at times *t* = 100, 200, 300 for three scenarios. The top row corresponds to a tumor in which both macrophage phenotypes are present and functional; the middle row depicts a scenario where only M_1_ macrophages are present (and alternative activation cannot occur), and the bottom row corresponds to a tumor whose macrophages are inert.

To quantify the dynamics of the cancer cells and macrophages, we calculate the average concentration, 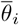, for *i* = *c, M*_1_, *M*_2_. 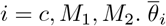 is calculated over the domain’s surface, *S*:

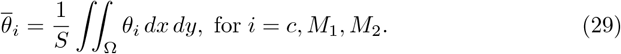

Figure 5 illustrates the average cancer cells concentration, 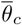, for the three afore-mentioned scenarios. If we view the scenario where macrophages are passive (case (iii)) as a reference point, we observe that the M_1_ phenotype significantly inhibits tumor expansion. However, M_1_ macrophages alone are unable to eradicate the tumor. By contrast, alternative activation and expression of the pro-tumor macrophage phenotype accelerate tumor growth.

**Fig. 5.**
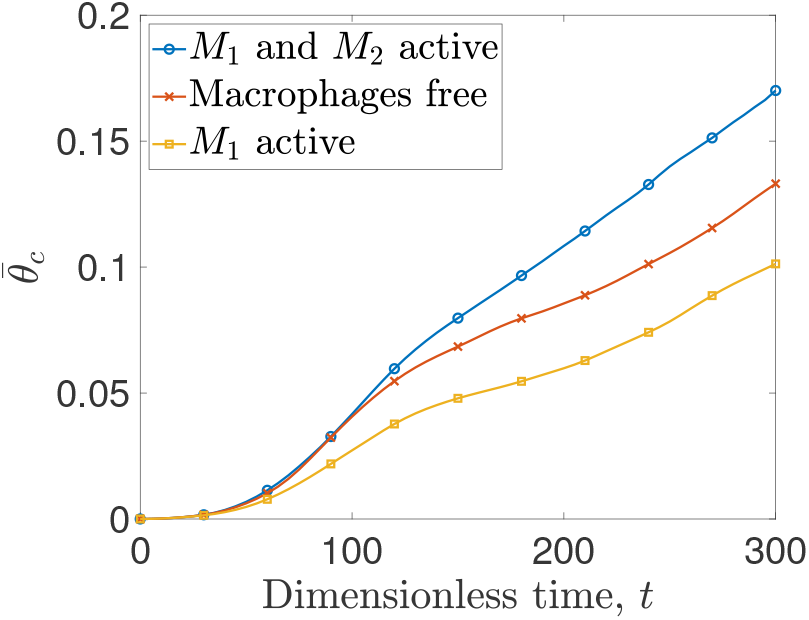
Average concentration, 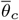, of cancer cells plotted against dimensionless time for three scenarios: a tumor with both macrophage phenotypes being expressed (blue curve with circles), macrophages being inert (red crossed curve), and macrophages immune to alternative activation (yellow curve with squares).

In Fig. 6 we plot the phase fluxes of the two macrophage phenotypes: 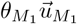, and 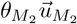, together with the spatial distribution of cancer cells at *t* = 200 and *t* = 300. The advancing tumor front and the forces it generates drive outward movement of both macrophage phenotypes. The fluxes for M_2_ macrophages (red arrows), are higher because chemotaxis drives them to regions with higher concentrations of CXCL12. The magnitude of the outward flux of M_1_ macrophages is smaller because they are subject to an inward chemotactic force, which drives them to hypoxic areas which are rich in CSF-1.

**Fig. 6.**
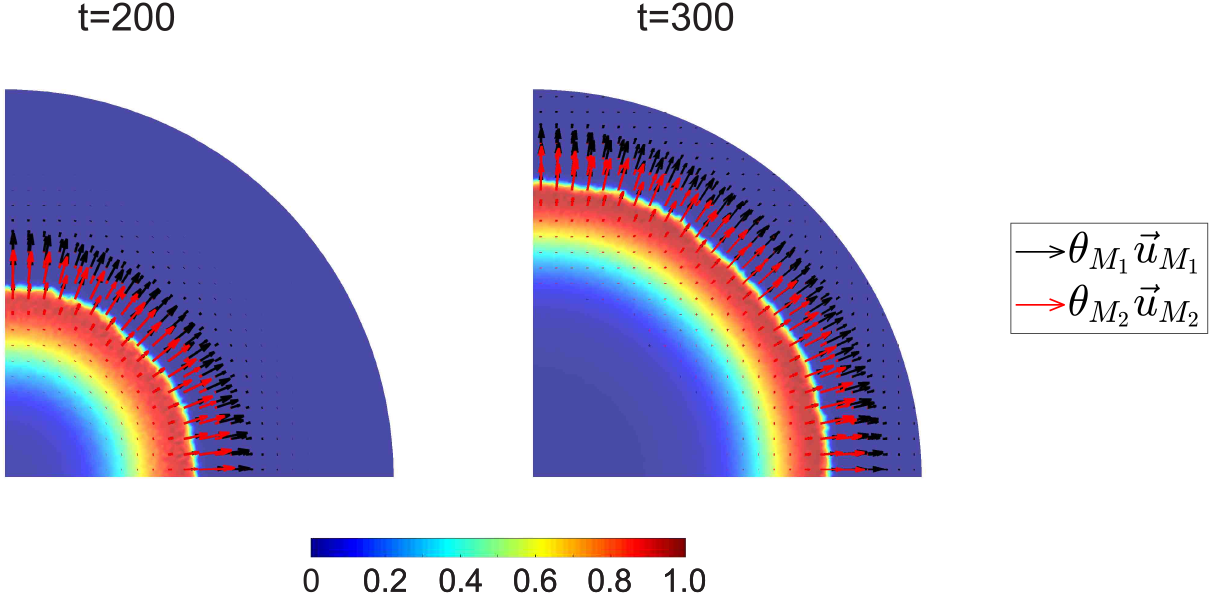
Phase flux vector distribution of macrophage phenotypes, alongside the surface distribution of cancer cells at *t* = 200 (left panel) and *t* = 300 (right panel). The *M*_1_ phase flux vectors are represented by red arrows, and the *M*_2_ phase flux vectors are depicted by black arrows.

### 4.0.1 Parametric analysis

We investigate the impact of varying some of the model parameters for which we lack accurate estimates. We focus on: (i) 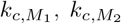 which represent the rates of interactions between cancer cells and *M*_1_, *M*_2_ macrophages, respectively (see Eq. (4)); (ii) *k*_*aa*_ which regulates the rate of alternative activation (see Eq. (8) and Eq. (9)); (iii) 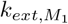 which denotes the *M*_1_ macrophage migration rate constant (see Eq. (8)); (iv) *χ*_*a*_, *χ*_*b*_ representing the chemosensitivity of M_1_ and M_2_ to spatial gradients of CSF-1 and CXCL12, respectively (see Eq. (1)). We quantify the impact of varying these parameters by either comparing their spatial distributions with the basic model or, recording the temporal evolution of the cancer cell average concentration, 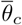.

Figure 7 (a) shows that increasing 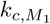 slows tumor growth, but does not lead to tumor eradication. In contrast, increasing 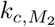 increases the tumor’s growth rate. Figure 7 (b) shows that changing *k*_*aa*_ by a factor of 5 significantly alters the tumor’s growth dynamics, with an increase in *k*_*aa*_ promoting tumor growth and a reduction inhibiting growth. However, further increasing or reducing this factor (to *×* 10 or *÷* 10) causes the corresponding curves to diverge less than one might anticipate. This suggests that there are limits to the attainable cancer cell average concentration, 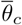, regardless of *k*_*aa*_ value; implying the existence of other limiting factors such as the rate of TGF-*β* replenishment in the tissue.

**Fig. 7.**
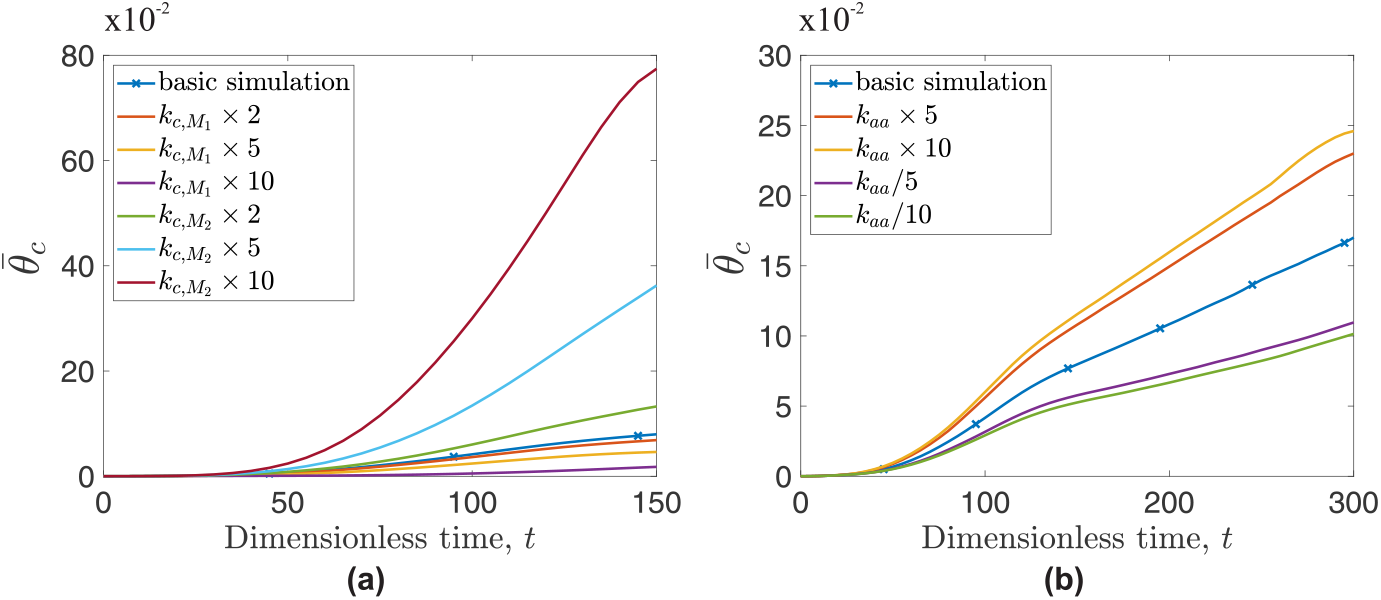
Temporal evolution of cancer cell average concentration, 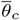 for simulations differing from the basic simulation by the value of one certain parameter: (a) 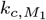 or 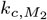 and (b) *k*_*aa*_. The parameter values for the basic simulation are: 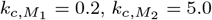, *k*_*aa*_ = 0.5.

We now take a closer look at some simulation results by examining the spatial distributions of cancer cells and macrophages at specific time points. Figure 8 compares the distributions of cancer cells and macrophages at *t* = 150 for the default case, with the distributions when *k*_*aa*_, 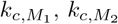, and 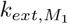 are increased by a factor of 5.

**Fig. 8.**
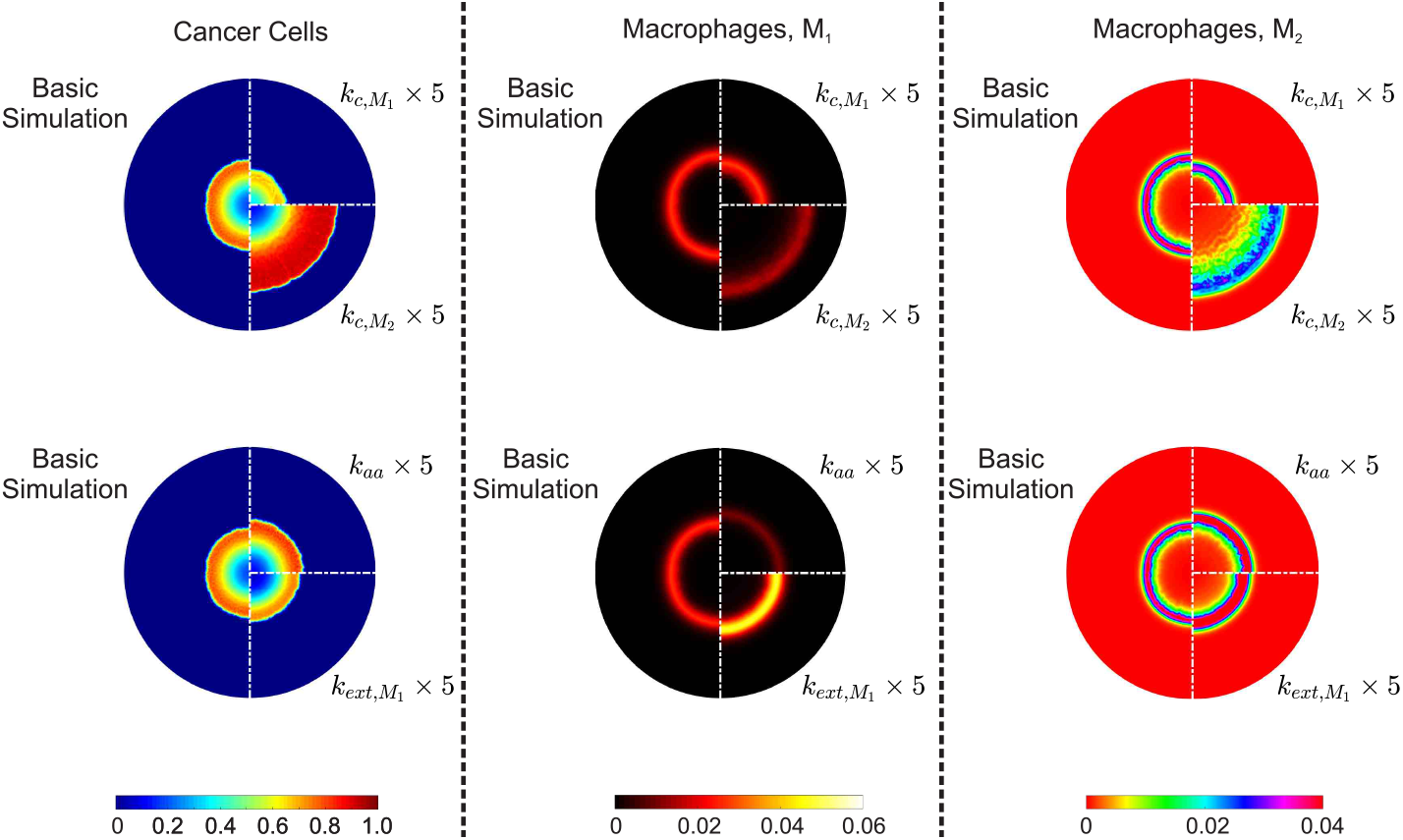
Surface distributions of cancer cells, *θ*_*c*_ (left column), macrophages 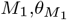 (middle column), and macrophages *M*_2_, 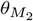 (right column) at dimensionless time, *t* = 150. The first row presents the impact of increasing the cancer cell-*M*_1_ interaction constant, 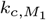, (top-right quadrant) and the cancer cell-*M*_2_ interaction constant, 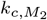 (bottom-right quadrant) by a factor of 5 compared to the basic simulation (left half-disk). The second row shows the impact of increasing the alternative activation rate constant, *k*_*aa*_ by a factor of 5 (top-right quadrant), and the effect of the migration rate constant, 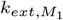 when increased by a factor of 5 (bottom-right quadrant).

Increasing the cancer cell-M_1_ interaction rate constant, 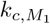, significantly reduces the cancer cell density and enables the immune cells to penetrate the tumor more effectively. The subsequent reduction in cancer cells leads to lower levels of cytokines, such as TGF-*β* and, hence, lower concentrations of *M*_2_ macrophages. However, the increased penetration of *M*_1_ macrophages results in a broader distribution of both macrophage phenotypes.

To highlight the differences between the system with enhanced cancer cell-*M*_2_ reaction rate constant, 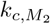, and the basic case, we use snapshots taken at *t* = 150 (see Fig. 8). The morphological differences between the two tumors are evident. The increased activity of pro-tumor macrophages leads to a tumor with a significantly wider and denser proliferating rim. Although the total number of immune cells in the tissue remains the same, they are distributed over a broader area due to the forces generated by the aggressive tumor growth.

Increasing *k*_*aa*_ produces a more aggressive tumor, with the proliferative rim showing higher cancer cell concentrations and the phenotypic balance shifting towards pro-tumor macrophages. Lastly, we enhance the migration rate constant, 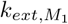; given that we are at *t* = 150, which is past the initial rapid accumulation of *M*_2_ macrophages, it is evident that the anti-tumor macrophages have begun to overwhelm the tumor, leading to tumor growth slowing down. Despite this, with the default parameter values (except for the increased 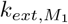) the anti-tumor macrophages remain largely confined to the tumor’s periphery. The influx of macrophages also naturally leads to an increase in pro-tumor macrophages (see Fig. 8).

#### 4.1 Multi-seed

In this section, we simulate tumors originating from different numbers of initial seeds, with both macrophage phenotypes present in their micro-environment. Specifically, in Fig. 9, we show how the spatialsurface distributions of cancer cells change over time for tissues seeded with one, two or three small tumors (columns 1, 2 and 3, respectively). Each simulation is initialised with the same number of tumor cells. Tumors originating from multiple seeds expand more rapidly than tumors initiated from a single lesion. Furthermore, hypoxic areas appear in the areas where the initial seeds merge.

**Fig. 9.**
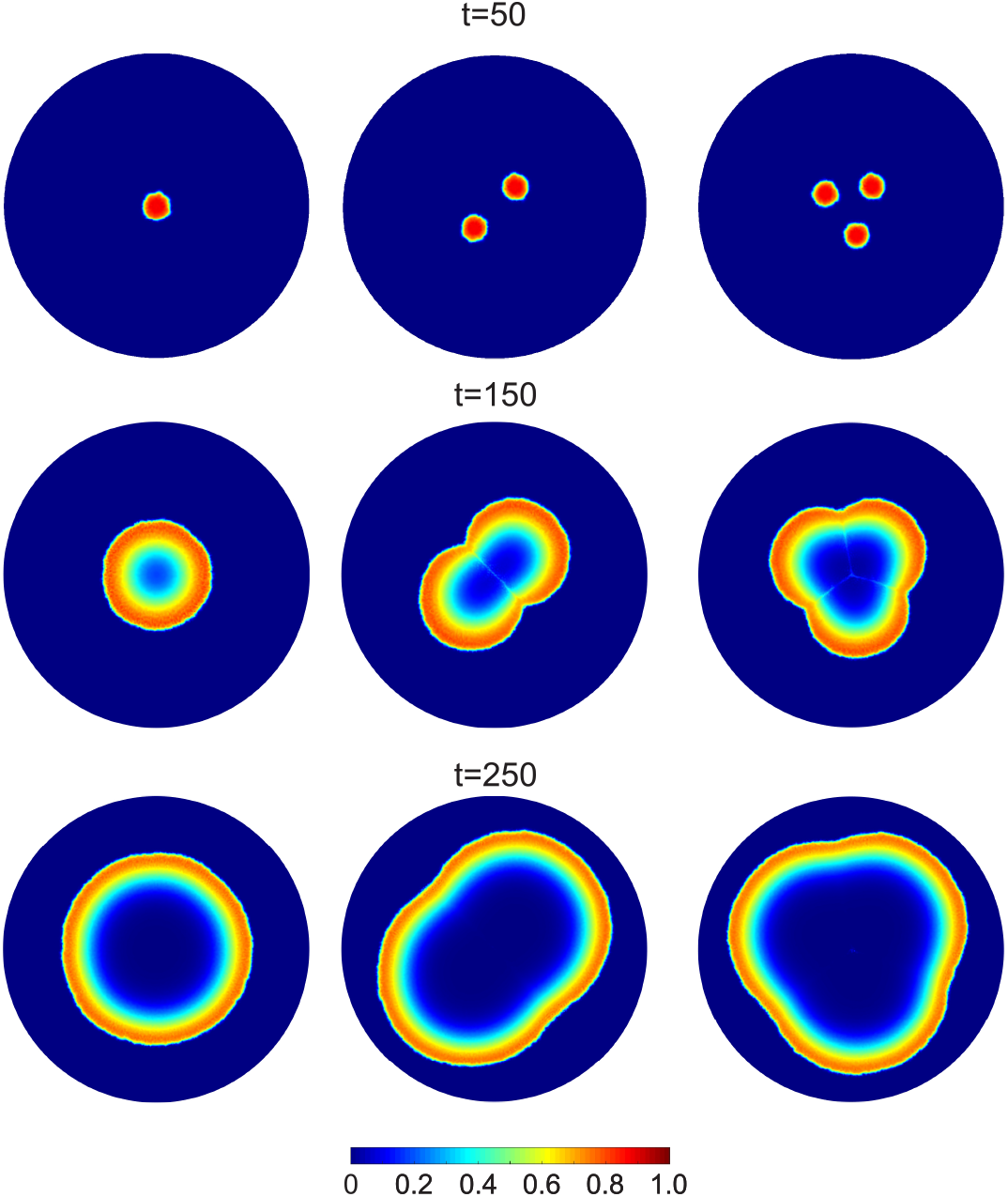
Spatial distributions of cancer cells for a tumor originating from a single (1st column), double (2nd column), and triple initial seed (3rd column) are depicted. At *t* = 0, the same number of cancer cells are divided equally between one, two or three seeds. Each row corresponds to a specific time point (*t* = 50, 150, and 250)

These zones become significant when we analyse the corresponding distributions of macrophages; Figure 10 shows that the pro-tumor macrophages are concentrated on the tumor’s periphery and the intersections formed between the two initial spheroids. In contrast to that, M_1_ macrophages are not present in this region.

**Fig. 10.**
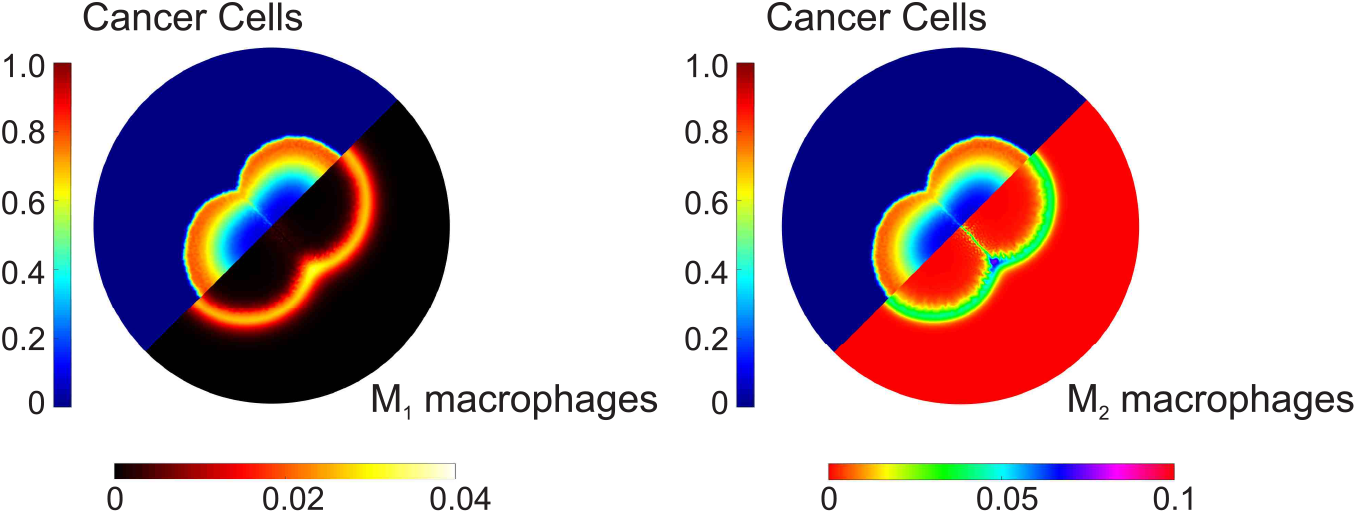
Juxtaposed spatial distributions of cancer cells with M_1_ macrophages (left panel) and cancer cells with M_2_ macrophages (right), for a tumor originating from a double seed at dimensionless time, *t* = 150. We observe a higher concentration of M_2_ macrophages at the intersection of the two tumor seeds.

M_2_ macrophages play a significant role in the tumor’s expansion, as evidenced in Fig. 11 (a), where we compare the cancer cell 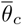 for tumors originating from single, double and triple seeds. While all three tumors initially contain the same number of cancer cells, distributing these cells into multiple seeds significantly enhances the tumor’s growth rate. This enhancement can be attributed partly to the higher perimeter-to-surface ratio resulting from distributing the same number of cells into more seeds, especially during the earlier stages of tumor development. An additional factor is the formation of hypoxic zones at the intersections of the seeds. These zones promote the activation of the M_2_ macrophage phenotype, thus contributing to tumor growth. The correlation between macrophage infiltration in hypoxic niches and a poorer prognosis is supported by experimental evidence [13].

**Fig. 11.**
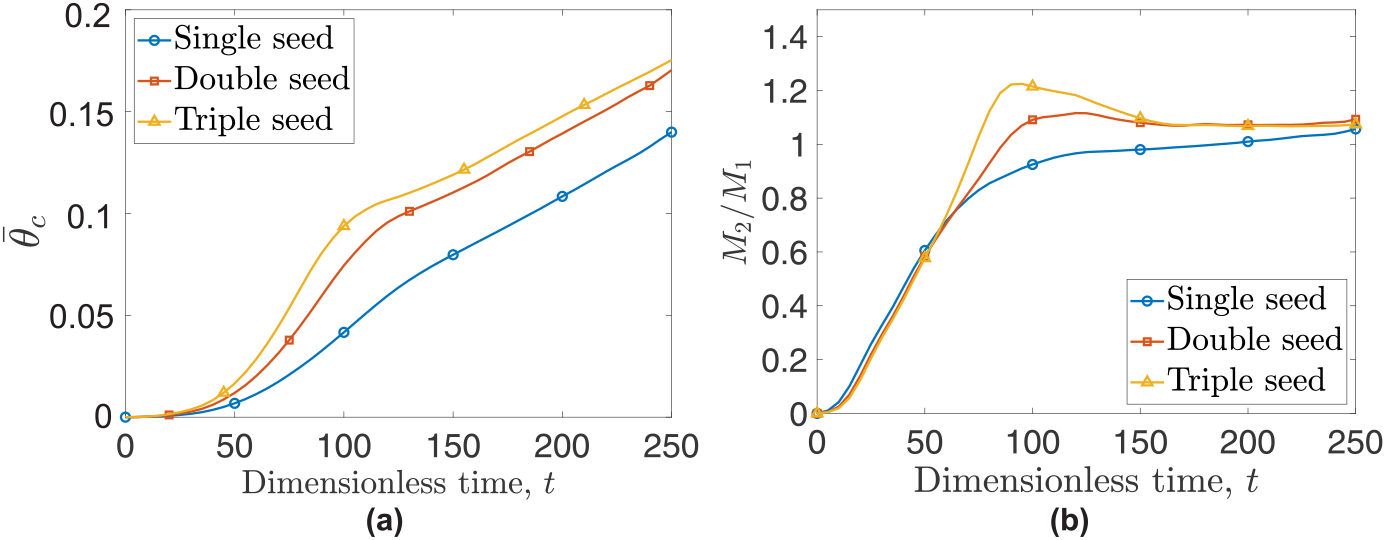
(a) Temporal evolution of cancer cell average concentration, 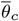, for tumors originating from one (blue curve with circles), two (red curve with rectangles), or three initial seeds (yellow curve with triangles). (b) Temporal evolution of the ratio of pro-tumor macrophages, M_2_, to pro-inflammatory macrophages, M_1_, for tumors originating from one, two or three initial seeds.

The formation of hypoxic pockets in tumors originating from multiple initial seeds also impacts the ratio of M_2_ to M_1_ macrophages, a metric generally associated with a poor prognosis [72]. Specifically, in Fig. 11 (b) we compare the time evolution of the M_2_ to M_1_ ratio for tumors origination from one, two or three seeds. The localized concentration of TAMs in hypoxic niches (observed in multi-seeds simulations) causes a spike in the M_2_/M_1_ ratio. This ratio can increase by up to 25% for a tumor originating from two seeds and up to 50% for a tumor originating from three seeds, when compared to a tumor originating from one seed. We also note that as the different tumor seeds merge to form a single mass, the *M*_2_*/M*_1_ ratio asymptotes to the same value regardless of the initial number of tumor seed.

## 5 Immunotherapy model

We now extend our model to investigate the efficacy of a macrophage targeting immunotherapy. We focus on the drug vactosertib, which binds to TGF-*β* receptors on M_1_ macrophages and, in doing so, inhibits their switching to a pro-tumor phenotype [48, 73]. In what follows, we will assume that M_1_ macrophages with bound vactosertib cannot change their phenotype. A graphical representation of the drug’s mechanism of action is presented in Fig. 12.

**Fig. 12.**
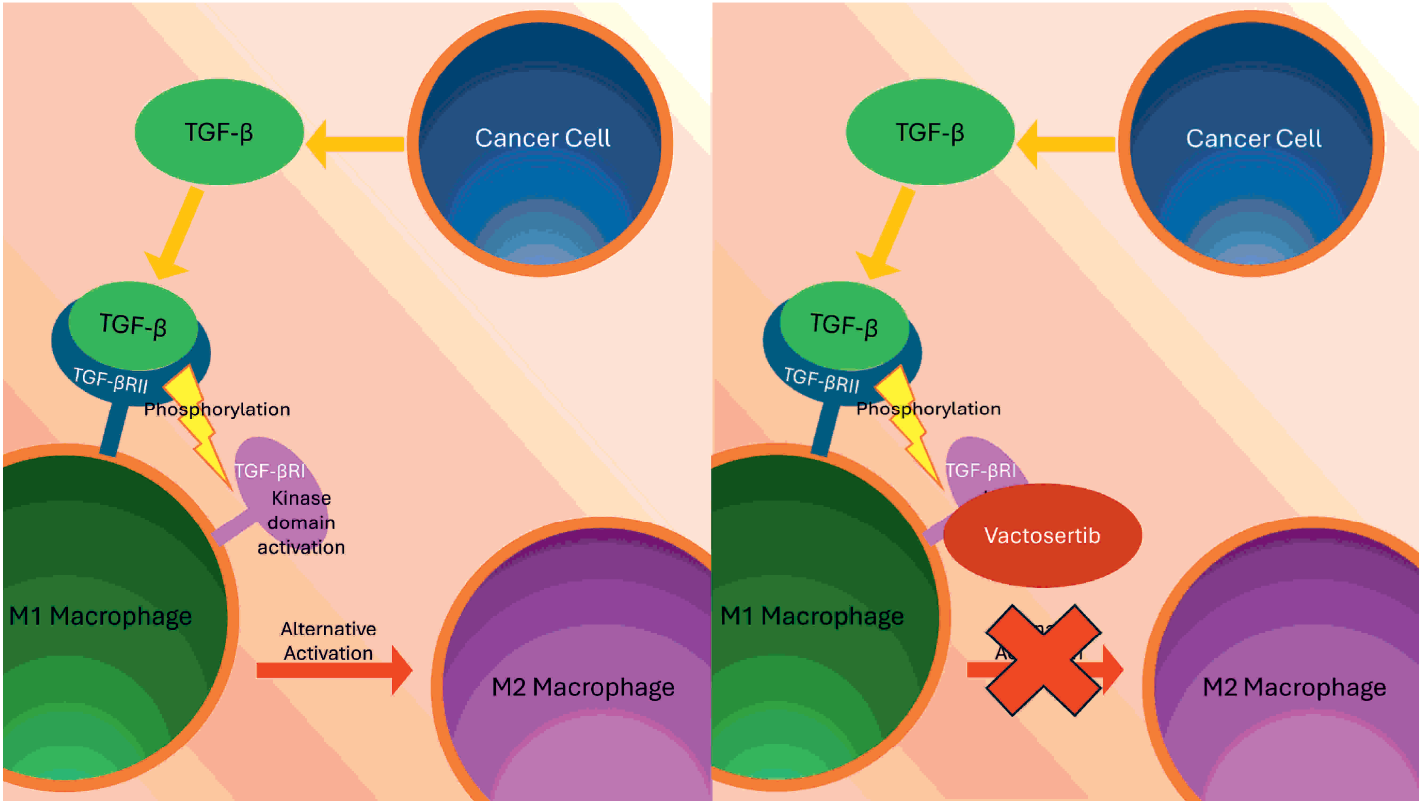
Schematic diagrams illustrating how vactosertib inhibits alternative activation of M_1_ macrophages. TGF-*β* produced by cancer cells binds to TGF-*β* type II receptors (TGF-*β*RII) on M_1_ macrophages, simulating the recruitment and phosphorylation of TGF-*β* type I receptors (TGF-*β*RI). This in turn activates the kinase domain of TGF-*β*RI [74]. In the presence of vactosertib, kinase activation on TGF-*β*RI is inhibited, preventing its phosphorylation and the subsequent alternative activation of the M_1_ macrophages [73].

In order to account for immunotherapy involving vactosertib, we include two new variables to our model. We denote by *d* the concentration of vactosertib and by 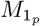 those M_1_ macrophages whose receptors are bound by the drug and which are, therefore, frozen in the M_1_ phenotype and resistant to alternative activation by TGF-*β*. We describe below the required model modifications.

### 5.1 Macrophages permanently exhibiting an anti-tumor phenotype

The mass balance equation for 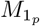 is similar to Eq. (1). In particular, the new sub-population exhibits the same phenotype and chemosensitivity as M_1_ macrophages.

Consequently, the mass balance equation for 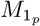 is:

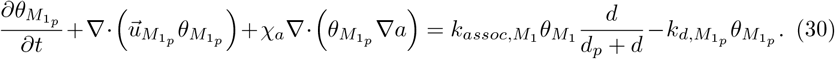

The source and sink terms on the right hand side of Eq. (30) represent the production of 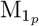 macrophages due to vactosertib binding to M_1_ macrophages and removal due to natural death. The parameter 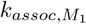 is the maximum rate of drug binding to the TGF-*β* receptors on the surface of M_1_ macrophages and *d*_*p*_ the drug concentration at which the association rate is half maximal. We assume that the vactosertib-M_1_ binding is irreversible. The sink term for 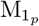 macrophages is identical to that for M_1_ macrophages (see Eq. (8)). 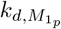 denotes the macrophages death rate constant (with 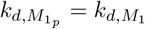).

The velocity field for 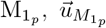, is calculated using Eq. (11), as described in Sec. 2.3. Including a new cellular phase affects the functional form of several other model equations. The 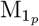 source term is balanced by a sink term for M_1_ macrophages:

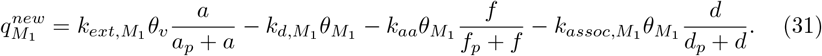

The following adjustments are made in the expressions for the source terms of cancer cells, *q*_*c*_, and vessels, *q*_*v*_:

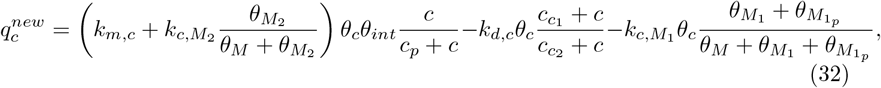

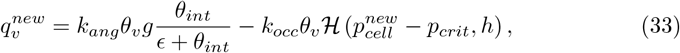

where

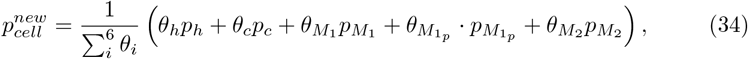

and 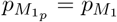.

### 5.2 Immunotherapeutic drug

In clinical trials vactosertib is administered orally twice a day [50] and reaches the tumor via the vasculature. Vactosertib’s mass balance follows the formulation presented in Eq. (19). The net production term *s*_*d*_ accounts for drug delivery, its removal due to binding to anti-tumor macrophage receptors, and removal due to natural decay:

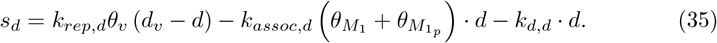

Here, *k*_*rep,d*_ denotes the drug delivery rate via the vasculature. *k*_*assoc,d*_ is the rate at which the drug binds irreversibly to TGF-*β* receptors on anti-tumor macrophages and, *k*_*d,d*_ denotes its rate of natural decay. We denote by *d*_*v*_ = *d*_*v*_ (*t*) the time-dependent drug concentration in the vasculature. The drug concentration in the vasculature takes its maximum value, *d*_*max*_, upon each oral administration and decays exponentially between doses:

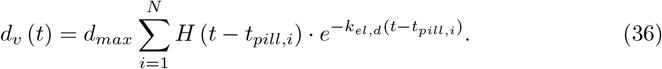

Here, *t*_*pill,i*_ is the time at which the *i*-th dose is delivered and

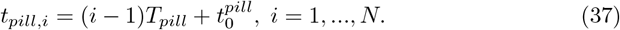

Where 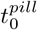 is the time at which treatment starts. *T*_*pill*_ is the time between doses and *N* is the number of doses administered. Additionally, *k*_*el,d*_ denotes the drug elimination rate within the vasculature and *H* denotes the Heaviside step function.

### 5.3 Parameters and parameter derivation

The parameters utilized in the immunotherapy model are presented in Table 6. All parameters are dimensionless (see Section II of the SI for details).

**Table 6.**
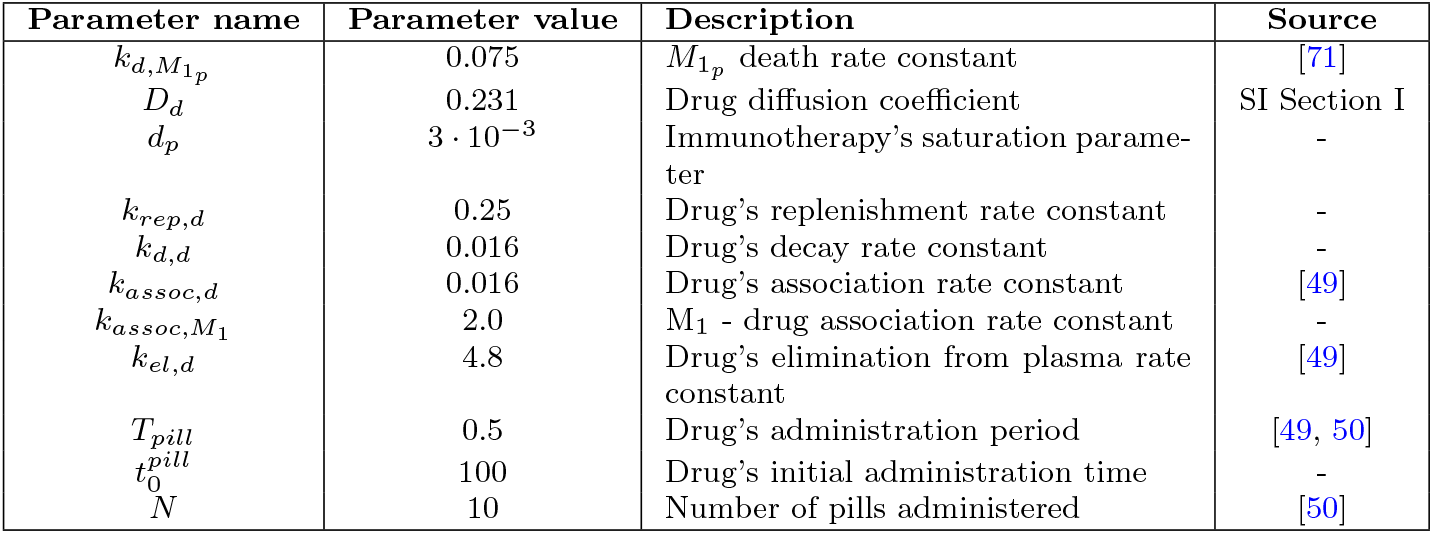
List of parameters associated with immunotherapy.

## 6 Immunotherapy model results

In this section, we present results from numerical simulations of our immunotherapy model. Vactosertib has shown promise in clinical trials [75–77], and additional clinical trials are currently underway [50–53].

For this computational study, we initiate immunotherapy at 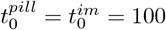 (see Eq. (36) and (37)). For comparison with the base case (Fig. 3), in Fig. 13 we plot the spatial distribution of cancer cells and macrophages at dimensionless times *t* = 150 and *t* = 250. We observe a significant delay in tumor growth (see distributions on left half of Fig. 13). The density of anti-tumor macrophages is larger when immunotherapy is applied, and the density of M_2_ macrophages is reduced. The anti-tumor macrophages increase in number and adopt a broader distribution, as the drug prevents them from switching to an *M*_2_ phenotype.

**Fig. 13.**
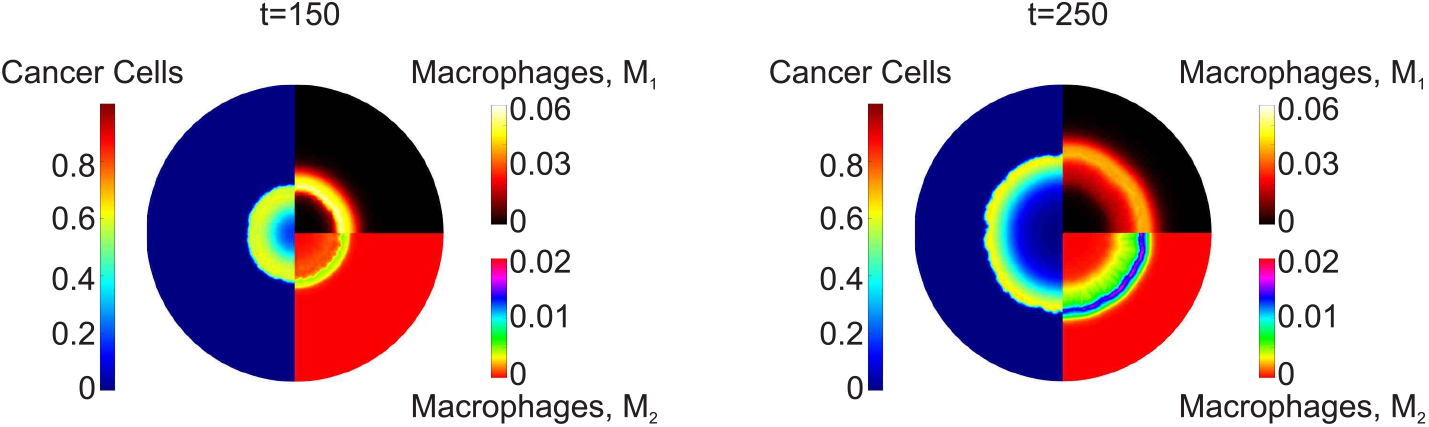
Surface distributions of cancer cells (left half-disk), *M*_1_ macrophages (upper-right quadrant), and *M*_2_ macrophages (bottom-right quadrant), at dimensionless times *t* = 150 (left column), and *t* = 250 (right column) for a tumor treated with immunotherapy. Immunotherapy is initiated at dimensionless time, 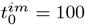.

Figure 14 (a) shows the impact of immunotherapy on the average concentration, 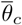, of cancer cells. Immunotherapy successfully halts cancer cell proliferation for a time interval of approximately Δ*t* ≈ 80 dimensionless time units (equivalent to a few months). At longer times, however, the tumor reverts to its initial growth rate due to the drug concentration’s decline in the tissue.

**Fig. 14.**
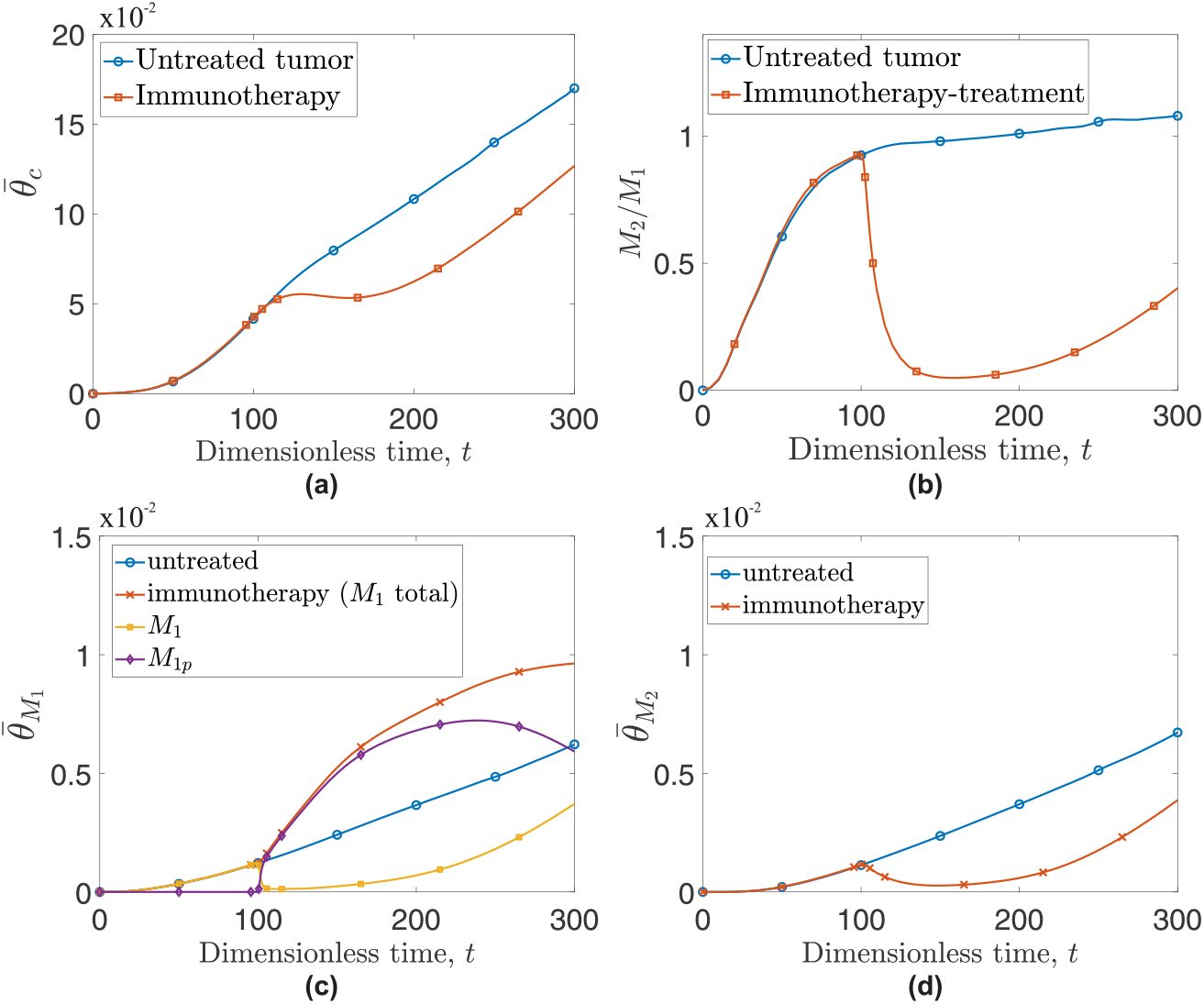
Evolution of (a) cancer cells average concentration, 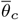, and of (b) the ratio of pro-tumor macrophages, M_2_ to pro-inflammatory macrophages, M_1_ for an untreated tumor (blue curve with circles) and a tumor treated with immunotherapy (red curve with squares). Evolution of average concentration of (c) M_1_ and (d) M_2_ macrophages for an untreated tumor (blue lines with circles) and an immunotherapy-treated tumor (red crossed lines). For the immunotherapy-treatment simulation, panel (c) also illustrates the evolution of drug-bound (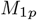 - purple line with diamonds) and unbound (M_1_ - yellow line with squares) macrophages.

Figure 14 (b) shows that immunotherapy significantly reduces the *M*_2_*/M*_1_ ratio by up to 95% for an extended time interval. However, as the drug’s effect begins to diminish (after a few months), the M_2_/M_1_ macrophage ratio increases and the tumor relapses. When calculating M_2_/M_1_ ratios, we use the sum of 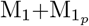 as a denominator. It is also of interest to examine the number of macrophages in the tissue. To achieve this, in Fig. 14 (c) and (d) we compare the dynamics of the 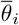 metrics for M_1_ and M_2_ macrophages for an untreated tumor (the basic model) with the dynamics for a tumor treated with vactosertib. Upon administration of the drug (at *t* = 100), we observe a rapid decline in the M_2_ phenotype, because the drug inhibits alternative activation of M_1_ macrophages. At the same time, being immune to alternative activation, M_1_ macrophages continue to rise in numbers, contributing to the sudden drop in the *M*_2_*/M*_1_ ratio observed in Fig. 14 (b). However, the circulatory system continues to serve as a pathway for newly recruited macrophages to enter the system. These macrophages continue to consume the administered drug and provide more receptors for TGF-*β*. Indeed, at dimensionless time, *t* ≈ 180, the population of *M*_2_ macrophages relapses and continues to increase thereafter. Gradual depletion of the drug slows the growth rate of drug-bound macrophages, and at dimensionless time *t* ≈ 235 we observe a decline in *M*_1*p*_. As the drug levels decline, the 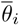 for both M_1_ and M_2_ macrophages adopt profiles similar to those observed in the untreated tumor.

In Fig. 15, we compare the average radial distributions of cancer cells and macrophages for the untreated and immunotherapy-treated tumors (rows 1 and 2 respectively) at dimensionless times, *t* = 200, and 300. For the treated tumor, we consider M_1_ macrophages as the sum of 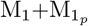 as both have anti-tumor properties. We observe that M_1_ macrophages, not only increase significantly in numbers in the presence of the immunotherapy drug but also gradually infiltrate the spheroid, resulting in a more uniform anti-tumor effect. Having said that, at *t* = 300, when the effect of the drug has largely dissipated, the profile of cancer cells looks similar for the treated and untreated cases.

**Fig. 15.**
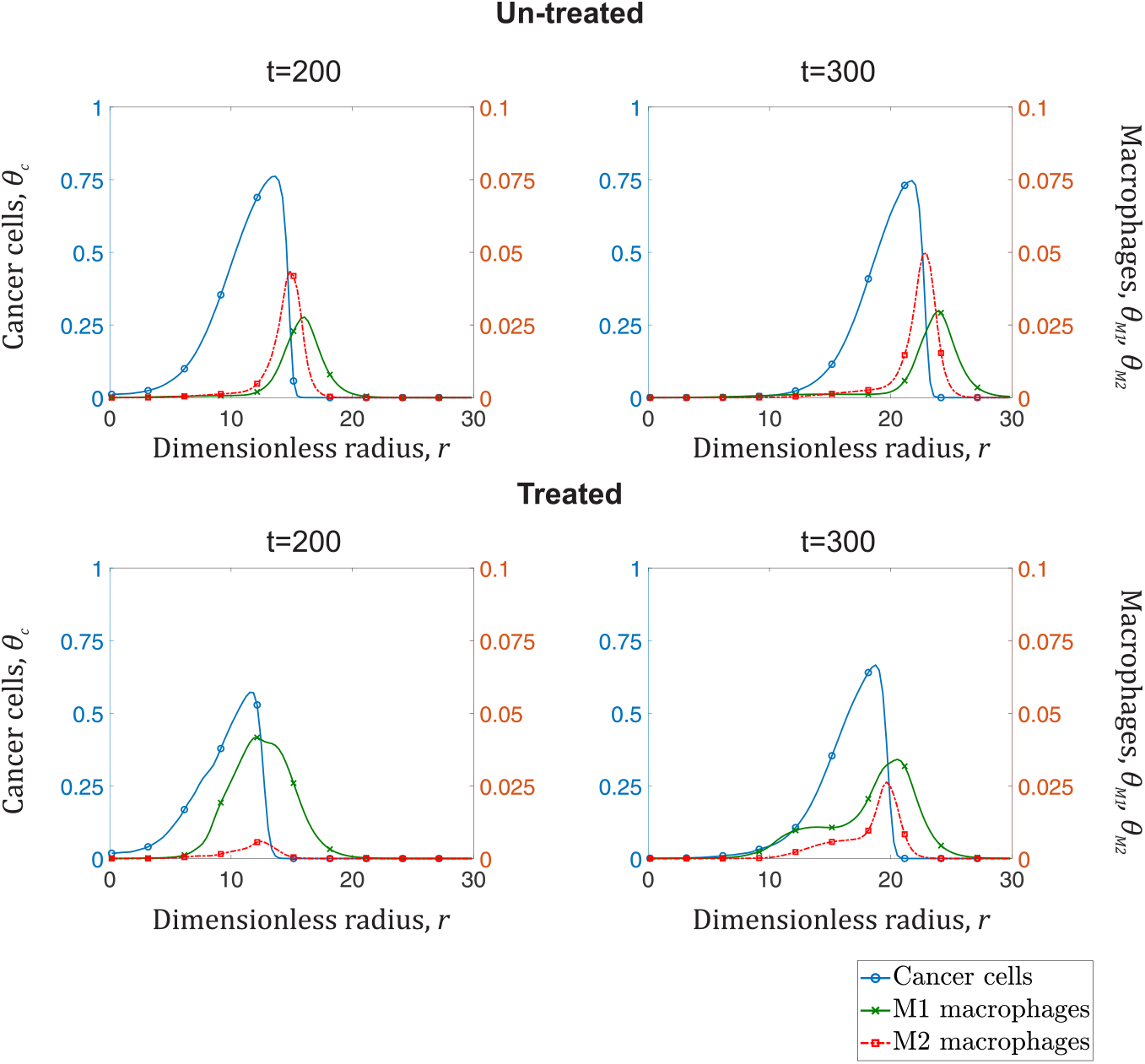
Average radial distributions of cancer cell concentrations (left y-axis - blue curve with open circles), M_1_ macrophage concentrations (right y-axis - green crossed curve), and M_2_ macrophage concentrations (right y-axis - red dotted curve with squares). Each column represents a specific time point (*t* = 200, and 300). Top row shows the radial distributions for an untreated tumor, and the bottom row depicts distributions for a tumor treated with immunotherapy.

### 6.1 Parametric analysis

In this section, we conduct a second parametric study, focusing initially on the parameters 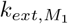 (macrophage migration rate constant), and *k*_*rep,d*_ (drug’s replenishment rate constant). Figure 16 shows how the dynamics of the cancer cell average concentration, 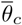, change as both parameters vary. In Fig. 16 (a), we observe that increasing 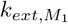 increases the cancer’s growth rate because the increased influx of M_1_ macrophages leads to a higher proportion of M_2_ macrophages. However, the effect is not significant. After the introduction of immunotherapy (*t* ≥ 100), higher 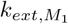 values cause a shift toward an anti-tumor *M*_1_ phenotype, driving a reduction in the tumor growth rate which becomes more pronounced as 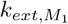 increases.

**Fig. 16.**
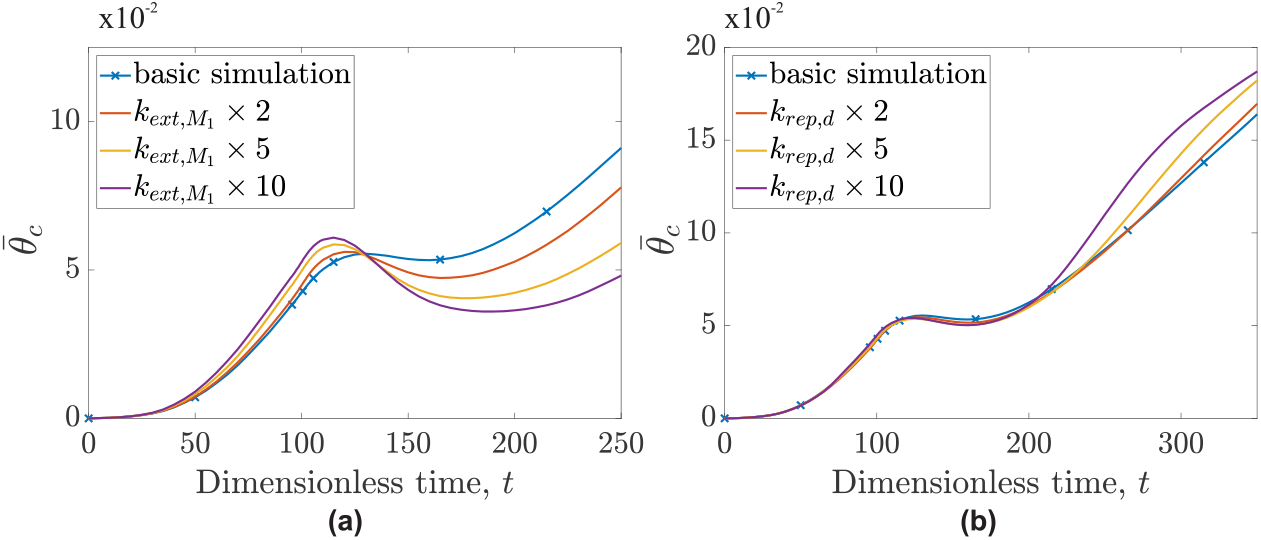
Temporal evolution of cancer cell average concentration, 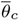, for scenarios differing from the basic immunotherapy (basic simulation) case by varying one parameter: (a) the macrophage migration rate constant, 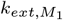 and (b) the drug replenishment rate, *k*_*rep,d*_.

Figure 16 (b) shows an interesting and unexpected result. Increasing the drug’s replenishment rate, *k*_*rep,d*_, while initially offering a modest short-term benefit, ultimately leads to a more aggressive tumor characterised by an increased growth rate. This behavior can be better understood by comparing the spatial distributions of the drug and TGF-*β*.

Figure 17 shows the average radial distributions of vactosertib and TGF-*β* for times 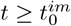. Two cases are considered: the basic simulation and one where *k*_*rep,d*_ is increased tenfold. Increasing *k*_*rep,d*_ increases the influx of vactosertib into the system. M_1_ macrophages present at that time, have their phenotype “frozen”. This increase in 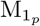 macrophages explains the modest reduction in the tumor’s growth rate. With the drug’s concentration remaining high and the influx of macrophages insufficient to bind it, vactosertib diffuses within the tumor, gradually moving behind the advancing TGF-*β* front formed by cancer cells, accumulating in an annulus extending from *r* ≈ 5 to *r* ≈ 10. These pockets of high drug concentration are further from new M_1_ macrophages than TGF-*β*. Thus, as new macrophages penetrate the tumor, they encounter TGF-*β* and adopt an M_2_ phenotype before they encounter the drug. Such behaviors are not observed for the default *k*_*rep,d*_. In the base-case, vactosertib is consumed by binding more gradually. Consequently, it diffuses less rapidly (due to the smaller spatial gradient) and does not accumulate behind the TGF-*β* front. These results explain why if *k*_*rep,d*_ values are large, immunotherapy may accelerate rather than slow tumor growth.

**Fig. 17.**
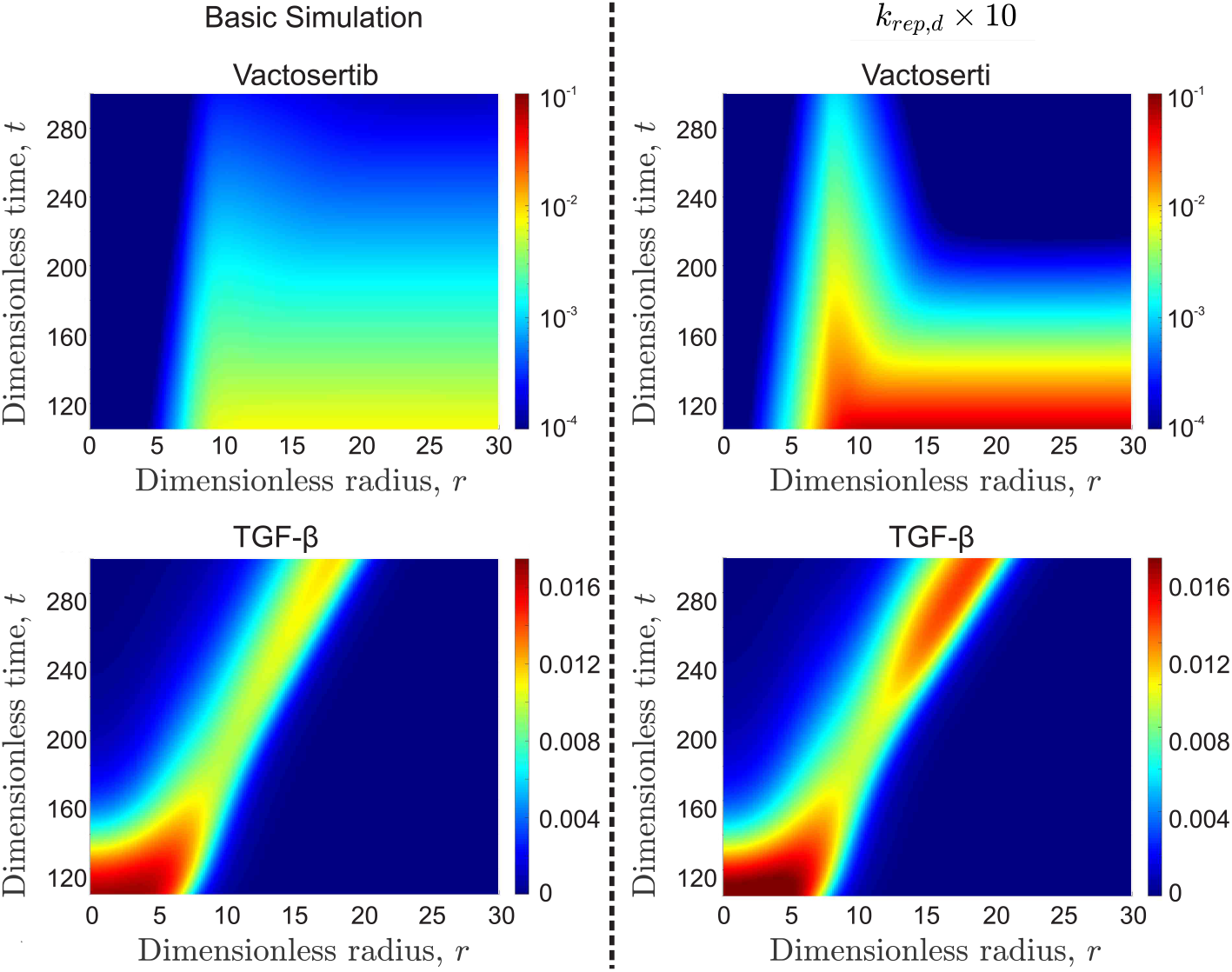
Evolution of averaged radial distributions of vactosertib (top row) and TGF-*β* (bottom row) for the base case (left column) and *k*_*rep,d*_ *×* 10 (right column).

## 7 Conclusions

In this study, we simulate the growth of a tumor embedded within healthy tissue and its infiltration by macrophages The model is based on mass and momentum balance equations applied to each cell phase (cellular species). To efficiently solve the model, we utilize the commercial software Comsol Multiphysics ^®^ and the Finite Elements Method (FEM).

The model captures a range of phenomena, particularly macrophage behavior, which is influenced by signals emitted by the tumor’s environment. To achieve this, the model takes into account chemical species closely related to the shaping of macrophage phenotype and activity.

The present model effectively captures the diverging behaviours of macrophages, distinguishing the pro-tumor action of M_2_ macrophages and the anti-cancer function of M_1_ macrophages. Furthermore, the model correlates the M_2_ to M_1_ ratio with the aggressiveness of tumor expansion and consequently, with a poorer expected outcome for the patient. This correlation is supported by experimental observations [72, 78]. Moreover, the model provides valuable insights into the macrophage distributions within the tumor’s micro-environment. Specifically, it accurately depicts macrophage localisation at the tumor’s periphery and the limited, yet noticeable, infiltration of macrophages into the tumor’s interior -a well documented behaviour [79, 80]. Moreover, a parametric study revealed the influence of certain parameters 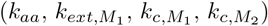 on the produced dynamics.

The spatial distributions of macrophages are further explored through simulations of multi-seed originating tumors. These simulations showcase the formation of hypoxic niches capable of accommodating dense populations of pro-tumor macrophages. These results are consistent with clinical images [79–81]. The simulation results show that in tumors originating from multi-seeds, M_2_ macrophages thrive within the hypoxic pockets formed when two spheroids meet, whereas M_1_ macrophages localise at the tumor periphery. This observation is consistent with experimental findings, in which TAMs found in hypoxic niches typically have the M_2_ phenotype, and the localisation of TAMS in such intersections has been reported previously [81]. Moreover, the infiltration of these cells into hypoxic niches is associated with a poorer prognosis for patients [13].

The study of multi-seed originating tumors and the integration of immunotherapy at a later stage serve as a testament to the model’s versatility. Indeed, the presented model effectively accommodates the description of radically different physics and seamlessly integrates therapy without imposing a significant increase in computational demands. The inclusion of immunotherapy is based on the compound vactosertib, which has demonstrated the ability to prevent the binding of TGF-*β* on macrophage receptors [48, 75] and is currently undergoing clinical trials as an anti-cancer treatment [50–53]. To the best of our knowledge, this is the first multiphase model to combine the study of macrophage behaviour, the phenomenon of alternative activation, and the administration of TGF-*β* receptor targeting immunotherapy.

The presented study offers ample opportunities for further research on tumor growth modeling. While the phenomenon of alternative activation is a central theme of the model, its counterpart, classical activation, remained unexplored. Investigating the conditions conducive to classical activation within the tumor’s environment presents a very intriguing topic with therapeutic implications. The present model serves as a natural foundation for a future comprehensive exploration of classical activation.

Additionally, in the present setting, only two populations of macrophages have been considered. Recent studies suggest [34, 35] the potential of consideration of multiple populations of macrophages and of continuous transitions between them. Similar approaches have been spearheaded by some of the present authors, e.g., in the context of the populations of stem cells [82, 83], and represent natural extensions of the model presented herein.

Regarding immunotherapy, the current model can accommodate other therapeutic factors. Macrophages represent an excellent target for a variety of immunotherapeutic compounds, several of which have undergone clinical trials (such as Lacnotuzumab [NCT02435680, NCT01643850] and Carlumab [NCT01204996]) [56]. It would be of particular interest to explore such combination therapies, since the mechanism considered herein based on vactosertib was not deemed sufficient (based on the mechanism considered herein) to lead to the full elimination of the cancer cell population. Moreover, the computational nature of the present study opens up the opportunity to alter the drug’s administration regimen. Although the biological feasibility of such a possibility merits further experimental studies, it is something we could explore computationally.

Lastly, another category of immune cells that play a significant role in cancer biology is T-cells. The ability of T-cells to identify and destroy cancer cells has generated considerable scientific interest, and the development of agents to enhance their anti-tumor responses holds great promise [84]. Such studies regarding the interplay of immune cells with other populations such as T-cells are presently of intense interest (see [85, 86] for recent examples), rendering the current setting a natural playground for related future studies.

## Supporting information

M1-M2_macrophages_SI

## Supplementary information

This article has one accompanying supplementary information file: “M1-M2 macrophages SI.pdf”.

## Data Availability

The data that support the findings of this study are available from the corresponding author upon reasonable request.

## References

[1] Hiam-Galvez, K.J., Allen, B.M., Spitzer, M.H.: Systemic immunity in cancer. Nature Reviews Cancer 21(6), 345–359 (2021)

[2] Gonzalez, H., Hagerling, C., Werb, Z.: Roles of the immune system in cancer: from tumor initiation to metastatic progression. Genes & Development 32(19-20), 1267–1284 (2018)

[3] Canadian Cancer Society: The immune system (2024). https://cancer.ca/en/cancer-information/what-is-cancer/immune-system

[4] Dunn, G.P., Old, L.J., Schreiber, R.D.: The three es of cancer immunoediting. Annual Review of Immunology 22(1), 329–360 (2004)

[5] Pittet, M.J., Nahrendorf, M., Swirski, F.K.: The journey from stem cell to macrophage. Annals of the New York Academy of Sciences 1319(1), 1–18 (2014)

[6] Perdiguero, E.G., Geissmann, F.: The development and maintenance of resident macrophages. Nature Immunology 17(1), 2–8 (2016)

[7] Ginhoux, F., Guilliams, M.: Tissue-resident macrophage ontogeny and homeostasis. Immunity 44(3), 439–449 (2016)

[8] Strell, C., Entschladen, F.: Extravasation of leukocytes in comparison to tumor cells. Cell Communication and Signaling 6, 1–13 (2008)

[9] Bied, M., Ho, W.W., Ginhoux, F., Blériot, C.: Roles of macrophages in tumor development: a spatiotemporal perspective. Cellular & Molecular Immunology 20(9), 983–992 (2023)

[10] Gao, J., Liang, Y., Wang, L.: Shaping polarization of tumor-associated macrophages in cancer immunotherapy. Frontiers in Immunology 13, 888713 (2022)

[11] Ross, E.A., Devitt, A., Johnson, J.R.: Macrophages: The good, the bad, and the gluttony. Frontiers in Immunology 12 (2021) 10.3389/fimmu.2021.708186

[12] Mosser, D.M., Edwards, J.P.: Exploring the full spectrum of macrophage activation. Nature Reviews Immunology 8(12), 958–969 (2008)

[13] Vito, A., El-Sayes, N., Mossman, K.: Hypoxia-driven immune escape in the tumor microenvironment. Cells 9(4), 992 (2020)

[14] Leung, E., Xue, A., Wang, Y., Rougerie, P., Sharma, V., Eddy, R., Cox, D., Condeelis, J.: Blood vessel endothelium-directed tumor cell streaming in breast tumors requires the hgf/c-met signaling pathway. Oncogene 36(19), 2680–2692 (2017)

[15] Wyckoff, J., Wang, W., Lin, E.Y., Wang, Y., Pixley, F., Stanley, E.R., Graf, T., Pollard, J.W., Segall, J., Condeelis, J.: A paracrine loop between tumor cells and macrophages is required for tumor cell migration in mammary tumors. Cancer Research 64(19), 7022–7029 (2004)

[16] Goswami, S., Sahai, E., Wyckoff, J.B., Cammer, M., Cox, D., Pixley, F.J., Stanley, E.R., Segall, J.E., Condeelis, J.S.: Macrophages promote the invasion of breast carcinoma cells via a colony-stimulating factor-1/epidermal growth factor paracrine loop. Cancer Research 65(12), 5278–5283 (2005)

[17] Arwert, E.N., Harney, A.S., Entenberg, D., Wang, Y., Sahai, E., Pollard, J.W., Condeelis, J.S.: A unidirectional transition from migratory to perivascular macrophage is required for tumor cell intravasation. Cell Reports 23(5), 1239–1248 (2018)

[18] Janssens, R., Struyf, S., Proost, P.: The unique structural and functional features of cxcl12. Cellular & Molecular Immunology 15(4), 299–311 (2018)

[19] Italiani, P., Töpfer, E., Boraschi, D.: Modulation of macrophage activation. In: Immune Rebalancing, pp. 123–149. Elsevier, ??? (2016)

[20] Zhang, F., Wang, H., Wang, X., Jiang, G., Liu, H., Zhang, G., Wang, H., Fang, R., Bu, X., Cai, S., et al.: Tgf-β induces m2-like macrophage polarization via snail-mediated suppression of a pro-inflammatory phenotype. Oncotarget 7(32), 52294 (2016)

[21] Darland, D.C., D’Amore, P.A., et al.: Blood vessel maturation: vascular development comes of age. The Journal of Clinical Investigation 103(2), 157–158 (1999)

[22] Gee, M.S., Procopio, W.N., Makonnen, S., Feldman, M.D., Yeilding, N.M., Lee, W.M.: Tumor vessel development and maturation impose limits on the effectiveness of anti-vascular therapy. The American Journal of Pathology 162(1), 183–193 (2003)

[23] Wu, W.-K., Llewellyn, O.P., Bates, D.O., Nicholson, L.B., Dick, A.D.: Il-10 regulation of macrophage vegf production is dependent on macrophage polarisation and hypoxia. Immunobiology 215(9-10), 796–803 (2010)

[24] Owen, M.R., Sherratt, J.A.: Pattern formation and spatiotemporal irregularity in a model for macrophage–tumour interactions. Journal of Theoretical Biology 189(1), 63–80 (1997)

[25] Owen, M.R., Sherratt, J.A.: Modelling the macrophage invasion of tumours: Effects on growth and composition. Mathematical Medicine and Biology: A Journal of the IMA 15(2), 165–185 (1998)

[26] Owen, M.R., Sherratt, J.A.: Mathematical modelling of macrophage dynamics in tumours. Mathematical Models and Methods in Applied Sciences 9(04), 513–539 (1999)

[27] Kelly, C., Leek, R., Byrne, H., Cox, S., Harris, A., Lewis, C.: Modelling macrophage infiltration into avascular tumours. Journal of Theoretical Medicine 4(1), 21–39 (2002)

[28] Owen, M.R., Byrne, H.M., Lewis, C.E.: Mathematical modelling of the use of macrophages as vehicles for drug delivery to hypoxic tumour sites. Journal of Theoretical Biology 226(4), 377–391 (2004)

[29] Webb, S.D., Owen, M.R., Byrne, H.M., Murdoch, C., Lewis, C.E.: Macrophage-based anti-cancer therapy: modelling different modes of tumour targeting. Bulletin of Mathematical Biology 69, 1747–1776 (2007)

[30] Leonard, F., Curtis, L.T., Yesantharao, P., Tanei, T., Alexander, J.F., Wu, M., Lowengrub, J., Liu, X., Ferrari, M., Yokoi, K., et al.: Enhanced performance of macrophage-encapsulated nanoparticle albumin-bound-paclitaxel in hypo-perfused cancer lesions. Nanoscale 8(25), 12544–12552 (2016)

[31] Boemo, M.A., Byrne, H.M.: Mathematical modelling of a hypoxia-regulated oncolytic virus delivered by tumour-associated macrophages. Journal of Theoretical Biology 461, 102–116 (2019)

[32] Louzoun, Y., Xue, C., Lesinski, G.B., Friedman, A.: A mathematical model for pancreatic cancer growth and treatments. Journal of Theoretical Biology 351, 74–82 (2014)

[33] Breems, N.Y., Eftimie, R.: The re-polarisation of m2 and m1 macrophages and its role on cancer outcomes. Journal of Theoretical Biology 390, 23–39 (2016)

[34] Eftimie, R.: Investigation into the role of macrophages heterogeneity on solid tumour aggregations. Mathematical Biosciences 322, 108325 (2020)

[35] Eftimie, R., Barelle, C.: Mathematical investigation of innate immune responses to lung cancer: The role of macrophages with mixed phenotypes. Journal of Theoretical Biology 524, 110739 (2021)

[36] Mahlbacher, G., Curtis, L.T., Lowengrub, J., Frieboes, H.B.: Mathematical modeling of tumor-associated macrophage interactions with the cancer microenvironment. Journal for Immunotherapy of Cancer 6, 1–17 (2018)

[37] Leonard, F., Curtis, L.T., Hamed, A.R., Zhang, C., Chau, E., Sieving, D., Godin, B., Frieboes, H.B.: Nonlinear response to cancer nanotherapy due to macrophage interactions revealed by mathematical modeling and evaluated in a murine model via crispr-modulated macrophage polarization. Cancer Immunology, Immunotherapy 69, 731–744 (2020)

[38] Suveges, S., Eftimie, R., Trucu, D.: Re-polarisation of macrophages within collective tumour cell migration: a multiscale moving boundary approach. Frontiers in Applied Mathematics and Statistics 7, 799650 (2022)

[39] Knútsdóttir, H., Pálsson, E., Edelstein-Keshet, L.: Mathematical model of macrophage-facilitated breast cancer cells invasion. Journal of Theoretical Biology 357, 184–199 (2014)

[40] Norton, K.-A., Jin, K., Popel, A.S.: Modeling triple-negative breast cancer heterogeneity: Effects of stromal macrophages, fibroblasts and tumor vasculature. Journal of Theoretical Biology 452, 56–68 (2018)

[41] Bull, J.A., Byrne, H.M.: Quantification of spatial and phenotypic heterogeneity in an agent-based model of tumour-macrophage interactions. PLOS Computational Biology 19(3), 1010994 (2023)

[42] Ward, J.P., King, J.R.: Mathematical modelling of avascular-tumour growth. Mathematical Medicine and Biology: A Journal of the IMA 14(1), 39–69 (1997)

[43] Byrne, H., Preziosi, L.: Modelling solid tumour growth using the theory of mixtures. Mathematical medicine and biology: a journal of the IMA 20(4), 341–366 (2003)

[44] Hubbard, M., Byrne, H.: Multiphase modelling of vascular tumour growth in two spatial dimensions. Journal of Theoretical Biology 316, 70–89 (2013)

[45] Breward, C.J., Byrne, H.M., Lewis, C.E.: A multiphase model describing vascular tumour growth. Bulletin of Mathematical Biology 65(4), 609–640 (2003)

[46] Breward, C., Byrne, H., Lewis, C.: The role of cell-cell interactions in a two-phase model for avascular tumour growth. Journal of Mathematical Biology 45(2), 125–152 (2002)

[47] Alard, E., Butnariu, A.-B., Grillo, M., Kirkham, C., Zinovkin, D.A., Newnham, L., Macciochi, J., Pranjol, M.Z.I.: Advances in anti-cancer immunotherapy: Cart cell, checkpoint inhibitors, dendritic cell vaccines, and oncolytic viruses, and emerging cellular and molecular targets. Cancers 12(7), 1826 (2020)

[48] Pubchem: Vactosertib Compound Summary (2024). https://pubchem.ncbi.nlm.nih.gov/compound/Vactosertib

[49] Jung, S.Y., Hwang, S., Clarke, J.M., Bauer, T.M., Keedy, V.L., Lee, H., Park, N., Kim, S.-J., Lee, J.I.: Pharmacokinetic characteristics of vactosertib, a new activin receptor-like kinase 5 inhibitor, in patients with advanced solid tumors in a first-in-human phase 1 study. Investigational New Drugs 38, 812–820 (2020)

[50] Samsung Medical Center (Responsible Party): Vactosertib With Nal-IRI/FL in Metastatic Pancreatic Ductal Adenocarcinoma (2024). https://clinicaltrials.gov/study/NCT04258072

[51] Case Comprehensive Cancer Center (Responsible Party): Oral TGF-beta Receptor I Inhibitor Vactosertib in SOC Chemoradiotherapy for Esophageal Adenocarcinoma (2024). https://clinicaltrials.gov/study/NCT06044311

[52] Yonsei University (Responsible Party): Antitumor Activity of Vactosertib in Combination With Pembrolizumab in Acral and Mucosal Melanoma Patients Progressed From Prior Immune Check Point Inhibitor. https://clinicaltrials.gov/study/NCT05436990

[53] Case Comprehensive Cancer Center (Responsible Party): Natural Killer (NK) Cells in Combination With Interleukin-2 (IL-2) and Transforming Growth Factor Beta (TGFbeta) Receptor I Inhibitor Vactosertib in Cancer. https://clinicaltrials.gov/study/NCT05400122

[54] Shojaee, P., Mornata, F., Deutsch, A., Locati, M., Hatzikirou, H.: The impact of tumor associated macrophages on tumor biology under the lens of mathematical modelling: A review. Frontiers in Immunology 13, 1050067 (2022)

[55] Mantovani, A., Allavena, P., Marchesi, F., Garlanda, C.: Macrophages as tools and targets in cancer therapy. Nature Reviews Drug Discovery 21(11), 799–820 (2022)

[56] Duan, Z., Luo, Y.: Targeting macrophages in cancer immunotherapy. Signal Transduction and Targeted Therapy 6(1), 127 (2021)

[57] Zhao, J., Huang, H., Zhao, J., Xiong, X., Zheng, S., Wei, X., Zhou, S.: A hybrid bacterium with tumor-associated macrophage polarization for enhanced photothermal-immunotherapy. Acta Pharmaceutica Sinica B 12(6), 2683–2694 (2022)

[58] Shields IV, C.W., Evans, M.A., Wang, L.L.-W., Baugh, N., Iyer, S., Wu, D., Zhao, Z., Pusuluri, A., Ukidve, A., Pan, D.C., et al.: Cellular backpacks for macrophage immunotherapy. Science Advances 6(18), 6579 (2020)

[59] Pyonteck, S.M., Akkari, L., Schuhmacher, A.J., Bowman, R.L., Sevenich, L., Quail, D.F., Olson, O.C., Quick, M.L., Huse, J.T., Teijeiro, V., et al.: Csf-1r inhibition alters macrophage polarization and blocks glioma progression. Nature Medicine 19(10), 1264–1272 (2013)

[60] Ries, C.H., Cannarile, M.A., Hoves, S., Benz, J., Wartha, K., Runza, V., Rey-Giraud, F., Pradel, L.P., Feuerhake, F., Klaman, I., et al.: Targeting tumorassociated macrophages with anti-csf-1r antibody reveals a strategy for cancer therapy. Cancer Cell 25(6), 846–859 (2014)

[61] Liu, X., Oh, S., Kirschner, M.W.: The uniformity and stability of cellular mass density in mammalian cell culture. Frontiers in Cell and Developmental Biology 10, 1017499 (2022)

[62] Harney, A.S., Arwert, E.N., Entenberg, D., Wang, Y., Guo, P., Qian, B.-Z., Oktay, M.H., Pollard, J.W., Jones, J.G., Condeelis, J.S.: Real-time imaging reveals local, transient vascular permeability, and tumor cell intravasation stimulated by tie2hi macrophage–derived vegfa. Cancer Discovery 5(9), 932–943 (2015)

[63] Wang, M., Yang, Y., Cansever, D., Wang, Y., Kantores, C., Messiaen, S., Moison, D., Livera, G., Chakarov, S., Weinberger, T., et al.: Two populations of self-maintaining monocyte-independent macrophages exist in adult epididymis and testis. Proceedings of the National Academy of Sciences 118(1), 2013686117 (2021)

[64] Mantovani, A., Marchesi, F., Malesci, A., Laghi, L., Allavena, P.: Tumour-associated macrophages as treatment targets in oncology. Nature Reviews Clinical Oncology 14(7), 399–416 (2017)

[65] Lampropoulos, I., Charoupa, M., Kavousanakis, M.: Intra-tumor heterogeneity and its impact on cytotoxic therapy in a two-dimensional vascular tumor growth model. Chemical Engineering Science 259, 117792 (2022)

[66] Lampropoulos, I., Kavousanakis, M.: Application of combination chemotherapy in two dimensional tumor growth model with heterogeneous vasculature. Chemical Engineering Science, 118965 (2023)

[67] Bernard, S., Herzel, H.: Why do cells cycle with a 24 hour period? Genome Informatics 17(1), 72–79 (2006)

[68] Hinow, P., Gerlee, P., McCawley, L.J., Quaranta, V., Ciobanu, M., Wang, S., Graham, J.M., Ayati, B.P., Claridge, J., Swanson, K.R., Loveless, M., Anderson, A.R.A.: A spatial model of tumor-host interaction: application of chemotherapy. Mathematical Biosciences and Engineering: MBE 6(3), 521 (2009)

[69] Folkman, J.: Tumor angiogenesis. Cancer 3, 355–388 (1975)

[70] Norton, K.-A., Popel, A.S.: Effects of endothelial cell proliferation and migration rates in a computational model of sprouting angiogenesis. Scientific Reports 6(1), 1–10 (2016)

[71] Parihar, A., Eubank, T.D., Doseff, A.I.: Monocytes and macrophages regulate immunity through dynamic networks of survival and cell death. Journal of Innate Immunity 2(3), 204–215 (2010)

[72] Jayasingam, S.D., Citartan, M., Thang, T.H., Mat Zin, A.A., Ang, K.C., Ch’ng, E.S.: Evaluating the polarization of tumor-associated macrophages into m1 and m2 phenotypes in human cancer tissue: technicalities and challenges in routine clinical practice. Frontiers in Oncology 9, 1512 (2020)

[73] Kim, B.-G., Malek, E., Choi, S.H., Ignatz-Hoover, J.J., Driscoll, J.J.: Novel therapies emerging in oncology to target the tgf-β pathway. Journal of Hematology & Oncology 14, 1–20 (2021)

[74] Tzavlaki, K., Moustakas, A.: Tgf-β signaling. Biomolecules 10(3), 487 (2020)

[75] Malek, E., Rana, P.S., Swamydas, M., Daunov, M., Miyagi, M., Murphy, E., Ignatz-Hoover, J.J., Metheny, L., Jin, K.S., Driscoll, J.J.: Vactosertib, a novel tgf-β1 type i receptor kinase inhibitor, improves t-cell fitness: a single-arm, phase 1b trial in relapsed/refractory multiple myeloma. Research Square (2023)

[76] Lee, H.-J.: Recent advances in the development of tgf-β signaling inhibitors for anticancer therapy. Journal of Cancer Prevention 25(4), 213 (2020)

[77] Metropulos, A.E., Munshi, H.G., Principe, D.R.: The difficulty in translating the preclinical success of combined tgfβ and immune checkpoint inhibition to clinical trial. EBioMedicine 86 (2022)

[78] Zhang, M., He, Y., Sun, X., Li, Q., Wang, W., Zhao, A., Di, W.: A high m1/m2 ratio of tumor-associated macrophages is associated with extended survival in ovarian cancer patients. Journal of Ovarian Research 7, 1–16 (2014)

[79] Albert, N.L., Unterrainer, M., Fleischmann, D., Lindner, S., Vettermann, F., Brunegraf, A., Vomacka, L., Brendel, M., Wenter, V., Wetzel, C., et al.: Tspo pet for glioma imaging using the novel ligand 18 f-ge-180: first results in patients with glioblastoma. European Journal of Nuclear Medicine and Molecular Imaging 44, 2230–2238 (2017)

[80] Khurana, A., Chapelin, F., Xu, H., Acevedo, J.R., Molinolo, A., Nguyen, Q., Ahrens, E.T.: Visualization of macrophage recruitment in head and neck carcinoma model using fluorine-19 magnetic resonance imaging. Magnetic Resonance in Medicine 79(4), 1972–1980 (2018)

[81] Arlauckas, S.P., Garren, S.B., Garris, C.S., Kohler, R.H., Oh, J., Pittet, M.J., Weissleder, R.: Arg1 expression defines immunosuppressive subsets of tumor-associated macrophages. Theranostics 8(21), 5842 (2018)

[82] Celora, G.L., Byrne, H.M., Zois, C.E., Kevrekidis, P.G.: Phenotypic variation modulates the growth dynamics and response to radiotherapy of solid tumours under normoxia and hypoxia. Journal of Theoretical Biology 527, 110792 (2021) 10.1016/j.jtbi.2021.110792

[83] Celora, G.L., Byrne, H.M., Kevrekidis, P.G.: Spatio-temporal modelling of phenotypic heterogeneity in tumour tissues and its impact on radiotherapy treatment. Journal of Theoretical Biology 556, 111248 (2023) 10.1016/j.jtbi.2022.111248

[84] Ahmed, H., Mahmud, A.R., Faijanur-Rob-Siddiquee, M., Shahriar, A., Biswas, P., Shimul, M.E.K., Ahmed, S.Z., Ema, T.I., Rahman, N., Khan, M.A., et al.: Role of t cells in cancer immunotherapy: Opportunities and challenges. Cancer Pathogenesis and Therapy 1(02), 116–126 (2023)

[85] Mohammad Mirzaei, N., Hao, W., Shariyari, L.: Investigating the spatial interaction of immune cells in colon cancer. iScience 10, 106596 (2023)

[86] Robertson-Tessi, M., El-Kareh, A., Goriely, A.: A mathematical model of tumor– immune interactions. Journal of theoretical biology 294, 56–73 (2012)

