## Supplementary material for "Spatio-temporal dynamics of M_1_ and M_2_ macrophages in a multiphase model of tumor growth": M1-M2_macrophages_SI

#### Supplementary Information

I. Lampropoulos

*School of Chemical Engineering, National Technical University of Athens, Athens, Greece*

P. Kevrekidis

*Department of Mathematics and Statistics, University of Massachusetts Amherst, Amherst, Massachusetts*

C. Zois

*Department of Radiotherapy and Oncology, Democritus University of Thrace, Alexandroupolis, Greece*

H. Byrne

*Mathematical Institute, University of Oxford, Oxford, England and Ludwig Institute for Cancer Research, University of Oxford, Oxford, England*

M. Kavousanakis

*School of Chemical Engineering, National Technical University of Athens, Athens, Greece*

### CONTENTS

**I. Parameters derivation** 3

I.A. Basic model 4

I.B. Immunotherapy parameters 5

**II. Non-dimensionalised model** 7

II.A. Mass balances 7

II.B. Momentum balances 11

II.C. Auxiliary expressions 11

**III. Parametric analysis** 12

III.A. Parametric analysis for the immunotherapy model 14

**References** 16

#### I. PARAMETERS DERIVATION

We report here all the calculations performed for the derivation or estimation of the selected parameter values. A substantial portion of these parameters are adapted from Hubbard and Byrne's[1] model or derived from the models proposed by Lampropoulos et al.[2,3] and Bull and Byrne[4], as will be discussed below. For the remaining parameters, estimations were made based on available literature.

To determine the characteristic length of the studied domain, a few factors must be considered. Firstly, angiogenesis typically manifests when a tumor exceeds $1-2\,mm$ in diameter[5–7]. Furthermore, tumor spheroids typically develop necrotic zones when their diameter reaches approximately $4\,mm$[8]. Based on these factors, it is reasonable to set the characteristic length of the domain (the tumor seed's diameter at $t=0$) withing the range of $[100\,\mu m, 1000\,\mu m]$. Therefore, the selected characteristic scale was set to $L_o = 400\,\mu m$.

In addition to defining the spatial characteristic unit, we define the characteristic unit for time. As reported in Sec. 2.4, the unit selected for temporal non-dimensionalisation is the mitosis rate of healthy cells, denoted as $k_{m,h}$. Typically, the life cycle of a eukaryotic cell lasts approximately $24\,h$. Therefore, we define $k_{m,h} = \frac{1}{24\,h} = 1.157 \cdot 10^{-5}\,s^{-1}$.

The concentration of hemoglobin (Hgb) in adult humans ranged typically ranges from 12 to $18\,g/dL$ ($12-16\,g/dL$ for females and $14-18\,g/dL$ for males)[9]. Therefore, selecting $C_{Hgb} = 140\,g/L$ for our calculations falls within the acceptable range. Furthermore, each molecule of Hgb can bind up to four oxygen molecules[10]. Combining this information with the fact that oxygen content in blood is approximately $20\frac{mL\,O_2}{dL}$[11], we can calculate the value of the parameter $c_v$:

$$\left.\begin{array}{r}\left.\begin{array}{r} C_{Hgb} = 140\,\dfrac{g}{L} \\ M_{Hgb} = 64458\,\dfrac{g}{mol}\end{array}\right\} C_{Hgb} \approx 0.00217\dfrac{mol}{L} \\ 4\dfrac{mol\,O_2}{mol\,Hgb}\end{array}\right\} c_v \approx 8.7 \cdot 10^{-3}\,M. \tag{1}$$

Lastly, it has been previously calculated that $k_{rep} = 0.0088\,s^{-1}$[3].

#### I.A. Basic model

While the values for VEGF, denoted as $g$, were initially adopted from Lampropoulos and Kavousanakis[3], we decided to rescale it as a variable, this time using $g_v = 10^{-12}\,M$. This decision stems from considering VEGF's physiological range in tissue, which typically falls within $[0.59\,pM, 0.65\,pM]$[12]. Selecting a characteristic value close to the physiological range not only makes computational sense but also enhances the interpretability of the results. With this rationale, we set the initial value of VEGF in the system as:

$$g'(x,y,0) = \frac{g(x,y,0)}{g_v} = \frac{0.6 \cdot 10^{-12}\,M}{10^{-12}\,M} = 0.6.$$

Setting the value of $g'$ in the initial conditions necessitates considering the parameter $k^*_{p,g}$ as a variable. Solving it yieldsthe parameter's value: $k^*_{p,g} = 0.00959$.

To calculate the dimensionless diffusion coefficients, we utilized the values reported in literature for CSF-1 ($D_a = 160\,\mu m^2 s^{-1}$), CXCL12 ($D_b = 150 \cdot 10^{-6}\,\mu m^2 s^{-1}$), EGF ($D_l = 160\,\mu m^2 s^{-1}$), TGF-$\beta$ ($D_f = 21.3\,\mu m^2 s^{-1}$)[4] and oxygen ($D_c = 1.4 \cdot 10^{-5}\,cm^2 s^{-1}$)[3,13].

$$D^*_i = \frac{D_i}{k_{rep} L_o^2} \text{ for } i = a,b,l,f \Leftrightarrow \frac{D^*_i}{D^*_c} = \frac{\frac{D_i}{\cancel{K_{rep} L_o^2}}}{\frac{D_c}{\cancel{K_{rep} L_o^2}}} \overset{D^*_c=1}{\Longleftrightarrow} D^*_i = \frac{D_i}{D_c} \Leftrightarrow \begin{cases} D^*_a = 0.11, \\ D^*_b = 0.001, \\ D^*_l = 0.11, \\ D^*_f = 0.5. \end{cases} \tag{2}$$

The values of chemotactic parameters $\chi_i$, for $i = a,b,l,f$, were based on calculations for $\chi_g$. This parameter has been measured at $\chi_g = 2600\,\frac{cm^2}{sM}$[14], thus its non-dimensionalisation is:

$$\chi^*_g = \frac{\chi_g \cdot g_v}{k_{m,h} \cdot L_o^2} = \frac{2600\,\frac{cm^2}{sM} \cdot 10^{-12} M}{1.157 \cdot 10^{-5}\,s^{-1} \cdot 16 \cdot 10^{-4} cm^2} = 0.14. \tag{3}$$

Based on the calculations of $\chi^*_g$, and considering that $a,b,l,f$ are all non-dimensionalised through $g_v$, it is reasonable to assume that $\chi_i$, for $i = a,b,l,f$, are of the same order of magnitude, with $\chi_b$ being enhanced to account for the increased motility of the $M_2$ phenotype.

The production rate constants $k^*_{p,i}$, for $i = a,b,l,f$, are estimated based on $k^*_{p,g} = 0.00959$: $k^*_{p,i} = 0.00959$, for $i = a,l,f$. The production of CXCL12 occurs in cells adjacent to the vasculature[4], but not by endothelial cells themselves. Therefore, to decouple the rate at which CXCL12 is secreted in the system from the concentration of the vasculature while maintaining the spatial correlation between the two, $k^*_{p,b}$ was re-scaled using $\theta_v(x,y,0)$:

$$k^*_{p,b} = \frac{k^*_{p,g}}{\theta_v(x,y,0)} \approx 0.55. \tag{4}$$

The remaining rate constants related to each chemical species are $k_{d,i}$ and $k_{assoc,i}$ $(i = a,b,l,f)$; the former represents the decay rate constant and the latter represents the molecule-cell binding rate constant. For the decay rate constants, the dimensionless values are determined based on measurements reported in the literature ($k_{d,a} = 1.9 \cdot 10^{-4} s^{-1}$, $k_{d,b} = 2 \cdot 10^{-5} s^{-1}$, $k_{d,l} = 1.9 \cdot 10^{-4} s^{-1}$, $k_{d,f} = 0.23\, min^{-1}$)[15–17]:

$$k^*_{d,i} = \frac{k_{d,i}}{k_{rep}} \text{ for } i = a,b,l,f \Leftrightarrow \begin{cases} k^*_{d,a} = 0.0216, \\ k^*_{d,b} = 0.227 \cdot 10^{-2}, \\ k^*_{d,l} = 0.0216, \\ k^*_{d,f} = 0.436. \end{cases} \tag{5}$$

On the other hand, the association rate constants are assumed based on the value of $k_{assoc.g}$:

$$k^*_{assoc,i} \approx 7 \cdot 10^{-6} \text{ for } i = a,b,l,f. \tag{6}$$

The values for the macrophage death rate constant are defined based on measurements of macrophage life expectancies in the simulated conditions[18,19].

Lastly, the saturation parameters $\theta_M$, $a_p$ and $f_p$ are selected based on the values of the variables $\theta_{M_1}$, $\theta_{M_2}$, $a$ and $f$, over the course of a simulation.

#### I.B. Immunotherapy parameters

The newly introduced parameters pertain to the immunotherapeutic drug and the drug-bound macrophages, which are considered a separate sub-population in the model. Firstly, we will present the calculations performed for determining the parameters associated with the drug. The drug's diffusion coefficient is determined through its hydrodynamic radius (calculated equal to $r_d \approx 0.659\, nm$[20], based on vactosertib's molecular weight, $M_{vac} = 399.4\, \frac{g}{mol}$[21]), using the Stokes - Einstein equation:

$$D = \frac{k_B T}{6\pi\eta r} \Rightarrow \frac{D^*_d}{\cancel{D^*_c}^{\,1}} = \frac{\frac{\cancel{k_B T}}{\cancel{6\pi\eta} r_c}}{\frac{\cancel{k_B T}}{\cancel{6\pi\eta} r_d}} = \frac{r_c}{r_d} \Rightarrow D^*_d = 0.23. \tag{7}$$

The drug's elimination rate from the vasculature, as well as the drug's absorption rate, are based on measurements reported by Jung et al[22]:

$$k^*_{el,d} = \frac{k_{el,d}}{k_{m,h}} = \frac{\frac{1}{5\,h}}{\frac{1}{24\,h}} = 4.8, \tag{8}$$

$$k^*_{assoc,d} = \frac{k_{assoc,d}}{k_{rep}} = \frac{\frac{1}{2\,h}}{0.0088\,s^{-1}} = 0.016. \tag{9}$$

The calculation of the therapy's period, $T_{pill}$, is based on both the regimen administered in the experiments reported by Jung et al.[22], and the ongoing clinical trials[23]. The latter are also considered for the determination of the number of therapeutic administrations ($N = 10$). Lastly, we make the following assumptions based on the results reported by Jung et al[22]: $k^*_{assoc,M_1} = 2$, $k^*_{assoc,d} = 0.016$.

#### II. NON-DIMENSIONALISED MODEL

In this section, we present the process of non-dimensionalisation that leads to the finalised expressions based on which calculations are performed. In order to non-dimensionalise the system's equations, the variables are rescaled using the following characteristic quantities:

$$t' = k_{m,h} \cdot t \quad c' = \frac{c}{c_v} \quad l' = \frac{l}{g_v}$$

$$\vec{x_i}' = \frac{\vec{x_i}}{L_o} \quad g' = \frac{g}{g_v} \quad f' = \frac{f}{g_v}$$

$$\vec{u_i}' = \frac{\vec{u_i}}{k_{m,h} L_o} \quad a' = \frac{a}{g_v} \quad d' = \frac{d}{d_{max}}$$

$$p_i' = \frac{p_i}{\Lambda} \quad b' = \frac{b}{g_v}$$

The phase concentrations are non-dimensionalised by the sum of all fluid/cellular concentrations at $t = 0$. There is no influx of mass in the system for $t = 0$:

$$\sum_i q_i = k_{ext,M_1} \theta_v \frac{\cancel{a}^0}{a_p + \cancel{a}^0} = 0.$$

This implies that for $t = 0$, the system can be considered closed, and the concentrations $\theta_i\,(x, y, t = 0)$ are defined as volume fractions of a fluid of constant volume. Therefore, the concentrations $\theta_i$ are non-dimensionalised as follows:

$$\begin{cases} \theta_i' = \frac{\theta_i}{\sum_i^6 \theta_i (x,y,t=0)} \\ \sum_i^6 \theta_i\,(x, y, t = 0) = 1 \end{cases} \Leftrightarrow \theta_i' = \theta_i.$$

##### II.A. Mass balances

Firstly, we present the mass balance non-dimensionalisations for both the fluid/cellular phases and the chemical species. Healthy cells mass balance:

$$\begin{gathered} \frac{\partial \theta_h}{\partial t} + \nabla \cdot (\vec{u_h}\theta_h) = k_{m,h}\theta_h\theta_{int}\frac{c}{c_p + c} - k_{d,h}\theta_h\frac{c_{c_1} + c}{c_{c_2} + c} \overset{\times \frac{1}{k_{m,h}}}{\Longleftrightarrow} \\ \frac{1}{k_{m,h}}\frac{\partial \theta_h}{\partial t} + \frac{1}{k_{m,h}}\frac{L_o}{L_o}\nabla \cdot (\vec{u_h}\theta_h) = \frac{\cancel{k_{m,h}}}{\cancel{k_{m,h}}}\theta_h\theta_{int}\frac{c/c_v}{c_p/c_v + c/c_v} + \frac{k_{d,h}}{k_{m,h}}\theta_h\frac{c_{c1}/c_v + c/c_v}{c_{c2}/c_v + c/c_v} \Leftrightarrow \\ \frac{\partial \theta_h}{\partial t'} + \nabla' \cdot \left(\vec{u_h'}\theta_h\right) = \theta_h\theta_{int}\frac{c'}{c_p^* + c'} - k_{d,h}^*\theta_h\frac{c_{c_1}^* + c'}{c_{c_2}^* + c'}. \end{gathered} \tag{10}$$

Cancer cells mass balance:

$$
\begin{aligned}
&\frac{\partial \theta_c}{\partial t} + \nabla \cdot (\vec{u_c}\theta_c) + \chi_l \nabla \cdot (\theta_c \nabla l) = \\
&\left(k_{m,c} + k_{c,M_2}\frac{\theta_{M_2}}{\theta_M + \theta_{M_2}}\right)\theta_c\theta_{int}\frac{c}{c_p + c} - k_{d,c}\theta_c\frac{c_{c_1} + c}{c_{c_2} + c} - k_{c,}\theta_c\frac{\theta_{M_1}}{\theta_M + \theta_{M_1}} \xLeftrightarrow{\times\frac{1}{k_{m,h}}} \\
&\frac{1}{k_{m,h}}\frac{\partial \theta_c}{\partial t} + \frac{1}{k_{m,h}}\frac{L_o}{L_o}\nabla \cdot (\vec{u_c}\theta_c) + \frac{1}{k_{m,h}}\frac{L_o^2}{L_o^2}\chi_l\frac{g_v}{g_v}\nabla \cdot (\theta_c \nabla l) = \\
&\left(\frac{k_{m,c}}{k_{m,h}} + \frac{k_{c,M_2}}{k_{m,h}}\frac{\theta_{M_2}}{\theta_M + \theta_{M_2}}\right)\theta_c\theta_{int}\frac{c/c_v}{c_p/c_v + c/c_v} - \frac{k_{d,c}}{k_{m,h}}\theta_c\frac{c_{c_1}/c_v + c/c_v}{c_{c_2}/c_v + c/c_v} - \frac{k_{c,M_1}}{k_{m,h}}\theta_c\frac{\theta_{M_1}}{\theta_M + \theta_{M_1}} \Leftrightarrow \\
&\frac{\partial \theta_c}{\partial t'} + \nabla' \cdot \left(\vec{u_c}'\theta_c\right) + \chi_l^* \nabla' \cdot (\theta_c \nabla' l') = \\
&\left(k_{m,c}^* + k_{c,M_2}^*\frac{\theta_{M_2}}{\theta_M + \theta_{M_2}}\right)\theta_c\theta_{int}\frac{c'}{c_p^* + c'} - k_{d,c}^*\theta_c\frac{c_{c_1}^* + c'}{c_{c_2}^* + c'} - k_{c,M_1}^*\theta_c\frac{\theta_{M_1}}{\theta_M + \theta_{M_1}}
\end{aligned} \tag{11}
$$

Capillaries mass balance:

$$
\begin{aligned}
&\frac{\partial \theta_v}{\partial t} + \nabla \cdot (\vec{u_v}\theta_v) + \chi_g \nabla \cdot (\theta_v \nabla g) = k_{ang}\theta_v g\frac{\theta_{int}}{\epsilon + \theta_{int}} - k_{occ}\theta_v\mathcal{H}\left(p_{cell} - p_{crit}, h\right) \xLeftrightarrow{\times\frac{1}{k_{m,h}}} \\
&\frac{1}{k_{m,h}}\frac{\partial \theta_v}{\partial t} + \frac{1}{k_{m,h}}\frac{L_o}{L_o}\nabla \cdot (\vec{u_v}\theta_v) + \frac{1}{k_{m,h}}\frac{L_o^2}{L_o^2}g_v\chi_g\nabla \cdot \left(\theta_v\nabla\frac{g}{g_v}\right) = \frac{k_{ang}\cdot g_v}{k_{m,h}}\theta_v\frac{g}{g_v}\frac{\theta_{int}}{\epsilon + \theta_{int}} - \frac{k_{occ}}{k_{m,h}}\theta_v\frac{\Lambda}{\Lambda}\mathcal{H}\left(p_{cell} - p\right. \text{[illegible]} \\
&\frac{\partial \theta_v}{\partial t'} + \nabla' \cdot \left(\vec{u_v}'\theta_v\right) + \chi_g^*\nabla' \cdot (\theta_v \nabla' g') = k_{ang}^*\theta_v g'\frac{\theta_{int}}{\epsilon + \theta_{int}} - k_{occ}^*\theta_v\mathcal{H}\left(p_{cell}' - p_{crit}^*, h^*\right)
\end{aligned} \tag{12}
$$

$M_1$ mass balance:

$$
\begin{aligned}
&\frac{\partial \theta_{M_1}}{\partial t} + \nabla \cdot (\vec{u_{M_1}}\theta_{M_1}) + \chi_a \nabla \cdot (\theta_{M_1}\nabla a) = k_{ext,M_1}\theta_v\frac{a}{a_p + a} - k_{d,M_1}\theta_{M_1} - k_{aa}\theta_{M_1}\frac{f}{f_p + f} \xLeftrightarrow{\times\frac{1}{k_{m,h}}} \\
&\frac{1}{k_{m,h}}\frac{\partial \theta_{M_1}}{\partial t} + \frac{1}{k_{m,h}}\frac{L_o}{L_o}\nabla \cdot (\vec{u_{M_1}}\theta_{M_1}) + \frac{1}{k_{m,h}}\frac{L_o^2}{L_o^2}g_v\chi_a\nabla \cdot \left(\theta_{M_1}\nabla\frac{a}{g_v}\right) = \\
&\frac{k_{ext,M_1}}{k_{m,h}}\theta_v\frac{a/g_v}{a_p/g_v + a/g_v} - \frac{k_{d,M_1}}{k_{m,h}}\theta_{M_1} - \frac{k_{aa}}{k_{m,h}}\theta_{M_1}\frac{f/g_v}{f_p/g_v + f/g_v} \Leftrightarrow \\
&\frac{\partial \theta_{M_1}}{\partial t'} + \nabla' \cdot \left(\vec{u_{M_1}}'\theta_{M_1}\right) + \chi_a^*\nabla' \cdot (\theta_{M_1}\nabla' a') = k_{ext,M_1}^*\theta_v\frac{a'}{a_p^* + a'} - k_{d,M_1}^*\theta_{M_1} - k_{aa}^*\theta_{M_1}\frac{f'}{f_p^* + f'}
\end{aligned} \tag{13}
$$

$M_2$ mass balance:

$$
\begin{aligned}
&\frac{\partial \theta_{M_2}}{\partial t} + \nabla \cdot \left(\vec{u_{M_2}} \theta_{M_2}\right) + \chi_b \nabla \cdot \left(\theta_{M_2} \nabla b\right) = k_{aa} \theta_{M_1} \frac{f}{f_p + f} - k_{d,M_2} \theta_{M_2} \xLeftrightarrow{\times \frac{1}{k_{m,h}}} \\
&\frac{1}{k_{m,h}} \frac{\partial \theta_{M_2}}{\partial t} + \frac{1}{k_{m,h}} \frac{L_o}{L_o} \nabla \cdot \left(\vec{u_{M_2}} \theta_{M_2}\right) + \frac{1}{k_{m,h}} \frac{L_o^2}{L_o^2} g_v \chi_b \nabla \cdot \left(\theta_{M_2} \nabla \frac{b}{g_v}\right) = \\
&\frac{k_{aa}}{k_{m,h}} \theta_{M_1} \frac{f/g_v}{f_p/g_v + f/g_v} - \frac{k_{d,M_2}}{k_{m,h}} \theta_{M_2} \Leftrightarrow \\
&\frac{\partial \theta_{M_2}}{\partial t'} + \nabla' \cdot \left(\vec{u_{M_2}}' \theta_{M_2}\right) + \chi_b^* \nabla' \cdot \left(\theta_{M_2} \nabla' b'\right) = k_{aa}^* \theta_{M_1} \frac{f'}{f_p^* + f'} - k_{d,M_2}^* \theta_{M_2}
\end{aligned}
\tag{14}
$$

$M_{1_p}$ mass balance:

$$
\begin{aligned}
&\frac{\partial \theta_{M_{1_p}}}{\partial t} + \nabla \cdot \left(\vec{u_{M_{1_p}}} \theta_{M_{1_p}}\right) + \chi_a \nabla \cdot \left(\theta_{M_{1_p}} \nabla a\right) = k_{assoc,M_1} \theta_{M_1} \frac{d}{d_p + d} - k_{d,M_{1_p}} \theta_{M_{1_p}} \xLeftrightarrow{\times \frac{1}{k_{m,h}}} \\
&\frac{1}{k_{m,h}} \frac{\partial \theta_{M_{1_p}}}{\partial t} + \frac{1}{k_{m,h}} \frac{L_o}{L_o} \nabla \cdot \left(\vec{u_{M_{1_p}}} \theta_{M_{1_p}}\right) + \frac{1}{k_{m,h}} \frac{L_o^2}{L_o^2} g_v \chi_a \nabla \cdot \left(\theta_{M_{1_p}} \nabla \frac{a}{g_v}\right) = \\
&\frac{k_{assoc,M_1}}{k_{m,h}} \theta_{M_1} \frac{d/d_{max}}{d_p/d_{max} + d/d_{max}} - \frac{k_{d,M_{1_p}}}{k_{m,h}} \theta_{M_{1_p}} \Leftrightarrow \\
&\frac{\partial \theta_{M_{1_p}}}{\partial t'} + \nabla' \cdot \left(\vec{u_{M_{1_p}}}' \theta_{M_{1_p}}\right) + \chi_a^* \nabla' \cdot \left(\theta_{M_{1_p}} \nabla' a'\right) = k_{assoc,M_1}^* \theta_{M_1} \frac{d'}{d_p^* + d'} - k_{d,M_{1_p}}^* \theta_{M_{1_p}}
\end{aligned}
\tag{15}
$$

Oxygen mass balance:

$$
\begin{aligned}
&D_c \nabla^2 c + k_{rep} \theta_v \left(c_v - c\right) - c \sum_{i=h,c} k_{c,i} \theta_i - \theta_{int} \frac{c}{c_p + c} \sum_{i=h,c} k_{cm,i} \theta_i = 0 \xLeftrightarrow{\times \frac{1}{k_{rep} c_v}} \\
&D_c \frac{1}{k_{rep}} \frac{L_o^2}{L_o^2} \nabla^2 \frac{c}{c_v} + \frac{\cancel{k_{rep}}}{\cancel{k_{rep}}} \theta_v \frac{\left(c_v - c\right)}{c_v} - \frac{c}{c_v} \sum_{i=h,c} \frac{k_{c,i}}{k_{rep}} \theta_i - \theta_{int} \frac{c/c_v}{c_p/c_v + c/c_v} \sum_{i=h,c} \frac{k_{cm,i}}{k_{rep}} \theta_i = 0 \Leftrightarrow \\
&D_c^* \nabla^2 c' + \theta_v \left(1 - c'\right) - c' \sum_{i=h,c} k_{c,i}^* \theta_i - \theta_{int} \frac{c'}{c_p^* + c'} \sum_{i=h,c} k_{cm,i}^* \theta_i = 0
\end{aligned}
\tag{16}
$$

VEGF mass balance:

$$
\begin{aligned}
&D_g \nabla^2 g + k_{p,g} \left(\theta_h + \theta_c + \theta_{M_2}\right) \frac{c}{\left(c_a + c\right)^2} - k_{assoc,g} \theta_v g - k_{d,g} \cdot g = 0 \xLeftrightarrow{\times \frac{1}{k_{rep} g_v}} \\
&D_g \frac{1}{k_{rep}} \frac{L_o^2}{L_o^2} \nabla^2 \frac{g}{g_v} + \frac{k_{p,g}}{k_{rep} c_v g_v} \left(\theta_h + \theta_c + \theta_{M_2}\right) \frac{c/c_v}{\left(c_a/c_v + c/c_v\right)^2} - \frac{k_{assoc,g}}{k_{rep}} \theta_v \frac{g}{g_v} - \frac{k_{d,g}}{k_{rep}} \cdot \frac{g}{g_v} = 0 \Leftrightarrow \\
&D_g^* \nabla^2 g' + k_{p,g}^* \left(\theta_h + \theta_c + \theta_{M_2}\right) \frac{c'}{\left(c_a^* + c'\right)^2} - k_{assoc,g}^* \theta_v g' - k_{d,g}^* \cdot g' = 0
\end{aligned}
$$

$$(17)$$

CSF-1 mass balance:

$$\begin{aligned}
&D_a \nabla^2 a + k_{p,a} \theta_c \frac{c}{\left(c_a + c\right)^2} - k_{d,a} \cdot a - k_{assoc,a} \theta_{M_1} a = 0 \xLeftrightarrow{\times \frac{1}{k_{rep} g_v}} \\
&D_a \frac{1}{k_{rep}} \frac{L_o^2}{L_o^2} \nabla^2 \frac{a}{g_v} + \frac{k_{p,a}}{k_{rep} c_v g_v} \theta_c \frac{c/c_v}{\left(c_a/c_v + c/c_v\right)^2} - \frac{k_{d,a}}{k_{rep}} \cdot \frac{a}{g_v} - \frac{k_{assoc,a}}{k_{rep}} \theta_{M_1} \frac{a}{g_v} = 0 \Leftrightarrow \\
&D_a^* \nabla^2 a' + k_{p,a}^* \theta_c \frac{c'}{\left(c_a^* + c'\right)^2} - k_{d,a}^* \cdot a' - k_{assoc,a}^* \theta_{M_1} a' = 0
\end{aligned} \qquad (18)$$

CXCL12 mass balance:

$$\begin{aligned}
&D_b \nabla^2 b + k_{p,b} \theta_v - k_{d,b} \cdot b - k_{assoc,b} \theta_{M_2} b = 0 \xLeftrightarrow{\times \frac{1}{k_{rep} g_v}} \\
&D_b \frac{1}{k_{rep}} \frac{L_o^2}{L_o^2} \nabla^2 \frac{b}{g_v} + \frac{k_{p,b}}{k_{rep} g_v} \theta_v - \frac{k_{d,b}}{k_{rep}} \cdot \frac{b}{g_v} - \frac{k_{assoc,b}}{k_{rep}} \theta_{M_2} \frac{b}{g_v} = 0 \Leftrightarrow \\
&D_b^* \nabla^2 b' + k_{p,b}^* \theta_v - k_{d,b}^* \cdot b' - k_{assoc,b}^* \theta_{M_2} b' = 0
\end{aligned} \qquad (19)$$

EGF mass balance:

$$\begin{aligned}
&D_l \nabla^2 l + k_{p,l} \theta_{M_2} - k_{d,l} \cdot l - k_{assoc,l} \theta_c \cdot l = 0 \xLeftrightarrow{\times \frac{1}{k_{rep} g_v}} \\
&D_l \frac{1}{k_{rep}} \frac{L_o^2}{L_o^2} \nabla^2 \frac{l}{g_v} + \frac{k_{p,l}}{k_{rep} g_v} \theta_{M_2} - \frac{k_{d,l}}{k_{rep}} \cdot \frac{l}{g_v} - \frac{k_{assoc,l}}{k_{rep}} \theta_c \frac{l}{g_v} = 0 \Leftrightarrow \\
&D_l^* \nabla^2 l' + k_{p,l}^* \theta_{M_2} - k_{d,l}^* \cdot l' - k_{assoc,l}^* \theta_c \cdot l' = 0
\end{aligned} \qquad (20)$$

TGF-$\beta$ mass balance:

$$\begin{aligned}
&D_f \nabla^2 f + k_{p,f} \theta_c - k_{d,f} \cdot f - k_{assoc,f} \theta_{M_1} f = 0 \xLeftrightarrow{\times \frac{1}{k_{rep} g_v}} \\
&D_f \frac{1}{k_{rep}} \frac{L_o^2}{L_o^2} \nabla^2 \frac{f}{g_v} + \frac{k_{p,f}}{k_{rep} g_v} \theta_c - \frac{k_{d,f}}{k_{rep}} \cdot \frac{f}{g_v} - \frac{k_{assoc,f}}{k_{rep}} \theta_{M_1} \frac{f}{g_v} = 0 \Leftrightarrow \\
&D_f^* \nabla^2 f' + k_{p,f}^* \theta_c - k_{d,f}^* \cdot f' - k_{assoc,f}^* \theta_{M_1} f' = 0
\end{aligned} \qquad (21)$$

Drug's mass balance:

$$\begin{aligned}
&D_d \nabla^2 d + k_{rep,d} \theta_v \left(d_c - d\right) - k_{assoc,d} \theta_{M_1} \cdot d - k_{d,d} \cdot d = 0 \xLeftrightarrow{\times \frac{1}{k_{rep} d_{max}}} \\
&D_d \frac{1}{k_{rep}} \frac{L_o^2}{L_o^2} \nabla^2 \frac{d}{d_{max}} + \frac{k_{rep,d}}{k_{rep}} \theta_v \frac{\left(d_c - d\right)}{d_{max}} - \frac{k_{assoc,d}}{k_{rep}} \theta_{M_1} \cdot \frac{d}{d_{max}} - \frac{k_{d,d}}{k_{rep}} \cdot \frac{d}{d_{max}} = 0 \Leftrightarrow \\
&D_d^* \nabla^2 d' + k_{rep,d}^* \theta_v \left(d_c^* - d'\right) - k_{assoc,d}^* \theta_{M_1} \cdot d' - k_{d,d}^* \cdot d' = 0
\end{aligned} \qquad (22)$$

where:

$$d_c^* = \frac{\cancel{d_{max}}}{\cancel{d_{max}}} \sum_{i=1}^{N} H\left(t' - t_{pill,i}^*\right) \cdot e^{-\frac{k_{el,d}}{k_{m,h}}\left(k_{m,h} \cdot t - k_{m,h} \cdot t_{pill,i}\right)} = \sum_{i=1}^{N} H\left(t' - t_{pill,i}^*\right) \cdot e^{-k_{el,d}^*\left(t' - t_{pill,i}^*\right)}. \qquad (23)$$

#### II.B. Momentum balances

$$\sum_{j,j\neq i} d^*_{i,j}\theta_i\theta_j\left(\vec{u'}_j-\vec{u'}_i\right)$$
$$-\theta_i\nabla\cdot\left(\Lambda^* p'_i \boldsymbol{I}\right)+\nabla\cdot\left[\theta_i\left[\mu^*_i\left(\nabla\vec{u'}_i+\left(\nabla\vec{u'}_i\right)^T\right)-\frac{2}{3}\mu^*_i\left(\nabla\cdot\vec{u'}_i\right)\boldsymbol{I}\right]\right]=\vec{0}, \tag{24}$$

for $i,j=h,c,v,int,M_1,M_2,M_{1_p}$, with $d^*_{i,j}=\frac{d_{i,j}}{d_{h,c}}, \Lambda^*=\frac{\Lambda}{d_{h,c}k_{m,h}L_o^2}, \mu^*_i=\frac{\mu_i}{d_{h,c}L_o^2}$.

#### II.C. Auxiliary expressions

Equation of continuity:

$$\frac{\partial}{\partial t}\sum_i\theta_i+\sum_i\nabla\cdot\left(\theta_i\vec{u}_i\right)+\chi_l\nabla\cdot\left(\theta_c\nabla l\right)+\chi_g\nabla\cdot\left(\theta_v\nabla g\right)+\chi_a\nabla\cdot\left(\theta_{M_1}\nabla a\right)+\chi_b\nabla\cdot\left(\theta_{M_2}\nabla b\right)=k_{ext,M_1}\theta_v a \; \text{[illegible]}$$
$$\frac{1}{k_{m,h}}\left(\frac{\partial}{\partial t}\sum_i\theta_i+\sum_i\nabla\cdot\left(\theta_i\vec{u}_i\right)+\chi_l\nabla\cdot\left(\theta_c\nabla l\right)+\chi_g\nabla\cdot\left(\theta_v\nabla g\right)+\chi_a\nabla\cdot\left(\theta_{M_1}\nabla a\right)+\chi_b\nabla\cdot\left(\theta_{M_2}\nabla b\right)\right)=\text{[illegible]}$$
$$\frac{\partial}{\partial t'}\sum_i\theta_i+\sum_i\nabla'\cdot\left(\theta_i\vec{u}_i'\right)+\chi^*_l\nabla'\cdot\left(\theta_c\nabla' l'\right)+\chi^*_g\nabla'\cdot\left(\theta_v\nabla' g'\right)+\chi^*_a\nabla'\cdot\left(\theta_{M_1}\nabla' a'\right)+\chi^*_b\nabla'\cdot\left(\theta_{M_2}\nabla' b'\right)=\text{[illegible]} \tag{25}$$

Equations of state:

$$p_h=p_c=p_{M_1}=p_{M_2}=p_{int}+\Sigma\left(\theta_h+\theta_c+\theta_{M_1}+\theta_{M_2}\right)\overset{\times\frac{1}{\Lambda}}{\Longleftrightarrow}$$
$$\frac{p_h}{\Lambda}=\frac{p_c}{\Lambda}=\frac{p_{M_1}}{\Lambda}=\frac{p_{M_2}}{\Lambda}=\frac{p_{int}}{\Lambda}+\frac{1}{\Lambda}\begin{cases}\frac{\Lambda\left(\theta_h+\theta_c+\theta_{M_1}+\theta_{M_2}-\theta^*\right)}{\left(\theta_v+\theta_{int}\right)^2}, & \text{if } \theta_h+\theta_c+\theta_{M_1}+\theta_{M_2}\geq\theta^*\\ 0, & \text{otherwise.}\end{cases}\Leftrightarrow$$
$$p'_h=p'_c=p'_{M_1}=p'_{M_2}=p'_{int}+\begin{cases}\frac{\left(\theta_h+\theta_c+\theta_{M_1}+\theta_{M_2}-\theta^*\right)}{\left(\theta_v+\theta_{int}\right)^2}, & \text{if } \theta_h+\theta_c+\theta_{M_1}+\theta_{M_2}\geq\theta^*\\ 0, & \text{otherwise.}\end{cases} \tag{26}$$

#### III. PARAMETRIC ANALYSIS

In this section we present complementary results from the parametric analysis presented in the main manuscript. We begin with Fig. 1 (a) and $\chi_a$. Increasing this parameter has a substantial impact on tumor growth. Enhancing the penetrative tendency of $M_1$

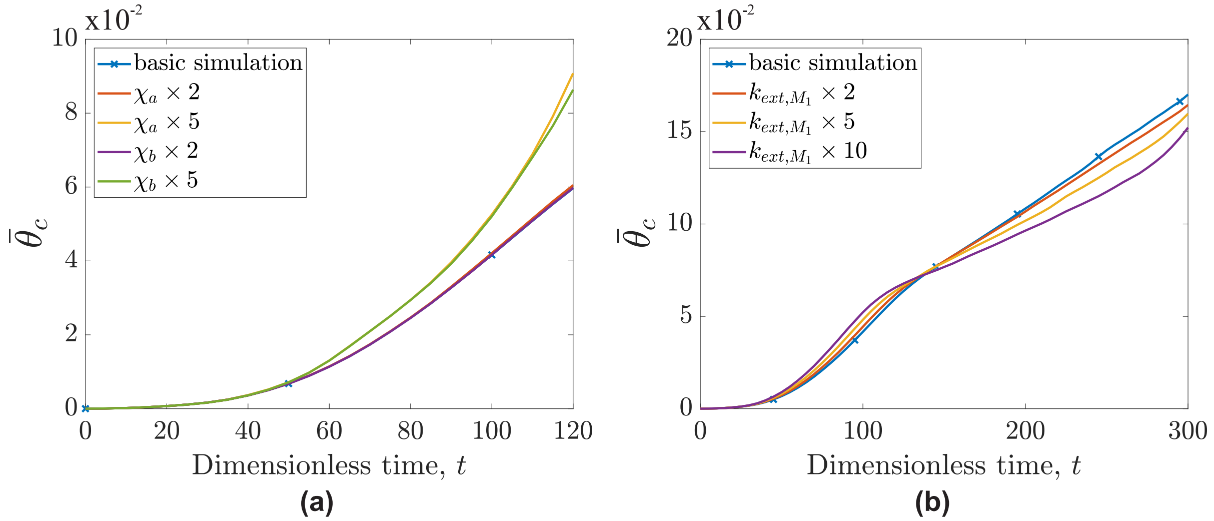


FIG. 1 Temporal evolution of cancer cell average concentration, $\overline{\theta}_c$, for parametric analysis of the parameters: (a) $\chi_a$ and $\chi_b$, and (b) $k_{ext,M_1}$.

macrophages increases the number of cells that reach the tumor's interior, where they are alternatively activated towards the $M_2$ phenotype. A similar effect is observed in Fig. 1 (a) with $\chi_b$. This time, it is the increased attraction of $M_2$ macrophages towards CXCL12 that allows them to reach the tumor's exterior, where they can more effectively exploit their pro-tumor phenotype more in a nutrient-rich environment and attract cancer cells toward it. In Fig. 1 (a) the presented times are cut short in comparison to the other figures. The reason behind that is the explosive nature of tumor expansion which makes computations severely harder. Lastly, Fig. 1 (b) shows the temporal evolution of $\overline{\theta}_c$ for tumors with different $k_{ext,M_1}$ values. This figure is particularly interesting as it suggests that an initially strong immune response can paradoxically assist the tumor by increasing the number of macrophages that are alternatively activated (towards $M_2$). However, beyond a certain point, the influx of new immune cells overwhelms the tumor's ability to influence them.

In Fig. 1 (b), we observe that all $\overline{\theta}_c$ curves converge around $t \approx 140$. To further investigate this behavior, Fig. 2 (a) presents the $\overline{\theta}_c$ evolution but this time with the alternative activation

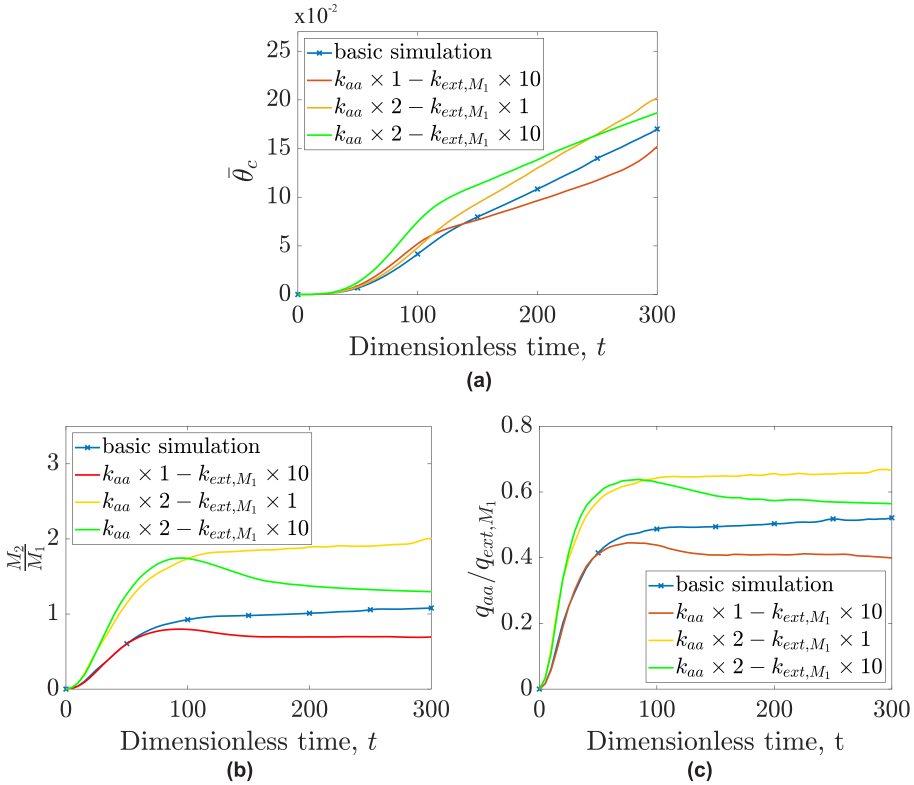


FIG. 2 (a) Temporal evolution of cancer cells average concentration, $\bar{\theta}_c$, with varying $k_{ext,M_1}$ and $k_{aa}$. (b) Temporal evolution of the total $M_2/M_1$ macrophages phenotype ratio for the same simulations. (c) Corresponding temporal evolution of the ratio of the alternative activation term, $q_{aa} \equiv k_{aa}\theta_{M_1}\frac{f}{f_p+f}$ (see Eq. (8) third term) to extravasation term, $q_{ext,M_1} \equiv k_{ext,M_1}\theta_v\frac{a}{a_p+a}$ (see Eq. (8) first term), for each calculation.

rate doubled. As expected, increasing the alternative activation rate accelerates tumor growth rate overall. What is particularly interesting is the emergence of a new convergence point between the two scenarios sharing the same $k_{aa}$ (one with the default $k_{ext,M_1}$ value and the other with 10 times that value). This convergence occurs later, at $t \approx 245$ suggesting that an enhanced ability to shift macrophages towards the pro-tumor state influences the system's behavior as $k_{ext,M_1}$ changes.

Figure 2 (b) supports these findings by presenting the $M_2/M_1$ ratio for the scenarios described. In cases with elevated $k_{aa}$, the resulting ratios are more than double those observed previously. However, by increasing the macrophage influx rate constant, $k_{ext,M_1}$, the relative contribution of $M_1$ macrophages steadily rises over time; this takes place as the system's capacity to convert incoming macrophages reaches its limit.

Figure 2 (c) provides further insight into the underlying mechanisms that produce these outcomes. The relative contributions and numbers of the two macrophage phenotypes are governed by two dominant processes: alternative activation and macrophage extravasation. When comparing the curves in Fig. 2 (c) with the macrophage ratios shown in Fig. 2 (b), it becomes apparent that they share many qualitative similarities. The equilibrium between these two processes ultimately determines the balance of macrophage phenotypes and significantly influences patient prognosis.

##### III.A. Parametric analysis for the immunotherapy model

This paragraph presents a complementary analysis for the parametric study of $k_{rep,d}$, conducted in the main manuscript. Figure 3 presents these distributions for: (a) $t = 150$ and default $k_{rep,d}$ values, (b) $t = 150$ and $k_{rep,d}$ increased tenfold, (c) $t = 250$ and default $k_{rep,d}$ values and, (b) $t = 250$ and $k_{rep,d}$ increased tenfold. The purpose of this figure is

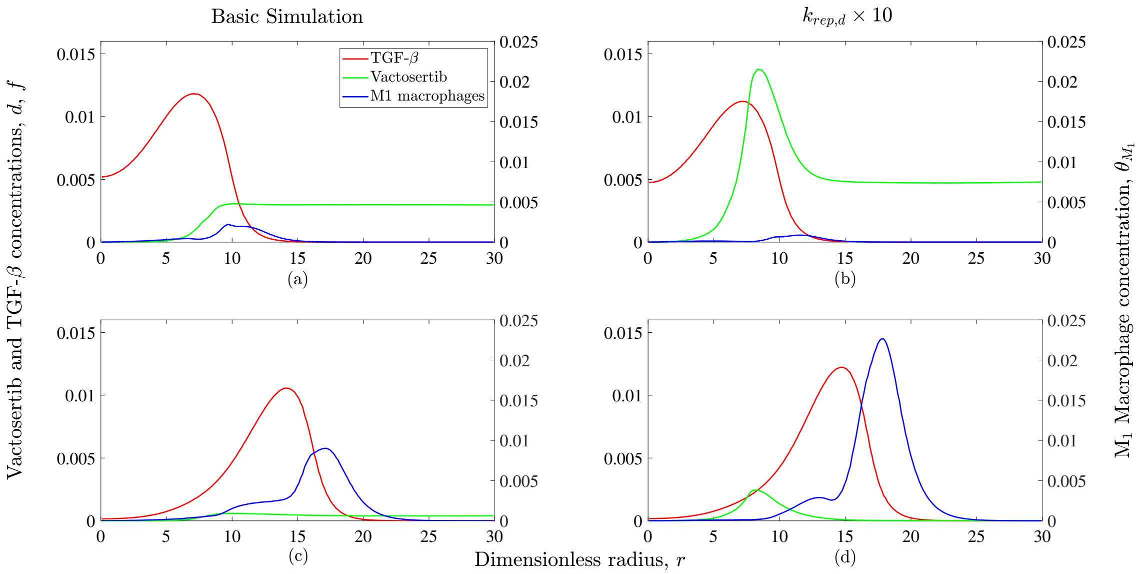


Fig. 3 Average radial distributions of $M_1$ macrophages, TGF-$\beta$ and, vactosertib for: (a) $t = 150$ and default $k_{rep,d}$, (b) $t = 150$ and increased $k_{rep,d}$, (c) $t = 250$ and default $k_{rep,d}$ and, (d) $t = 250$ and increased $k_{rep,d}$.

to present the distribution and abundance of vactosertib in relation to TGF-$\beta$ and -most importantly- $M_1$ macrophages. Here, we can better observe how increased vactosertib influx results in almost all macrophages having their phenotype frozen early on, leading to new macrophages gradually being separated from the drug's supplus due to the wall of TGF-$\beta$ that is located between them.

### REFERENCES


[1]M. Hubbard and H. Byrne, "Multiphase modelling of vascular tumour growth in two spatial dimensions," Journal of Theoretical Biology **316**, 70–89 (2013).

[2]I. Lampropoulos, M. Charoupa, and M. Kavousanakis, "Intra-tumor heterogeneity and its impact on cytotoxic therapy in a two-dimensional vascular tumor growth model," Chemical Engineering Science **259**, 117792 (2022).

[3]I. Lampropoulos and M. Kavousanakis, "Application of combination chemotherapy in two dimensional tumor growth model with heterogeneous vasculature," Chemical Engineering Science , 118965 (2023).

[4]J. A. Bull and H. M. Byrne, "Quantification of spatial and phenotypic heterogeneity in an agent-based model of tumour-macrophage interactions," PLOS Computational Biology **19**, e1010994 (2023).

[5]P. Hinow, P. Gerlee, L. J. McCawley, V. Quaranta, M. Ciobanu, S. Wang, J. M. Graham, B. P. Ayati, J. Claridge, K. R. Swanson, M. Loveless, and A. R. A. Anderson, "A spatial model of tumor-host interaction: application of chemotherapy," Mathematical Biosciences and Engineering: MBE **6**, 521 (2009).

[6]J. Folkman, "Tumor angiogenesis," Cancer **3**, 355–388 (1975).

[7]K.-A. Norton and A. S. Popel, "Effects of endothelial cell proliferation and migration rates in a computational model of sprouting angiogenesis," Scientific Reports **6**, 1–10 (2016).

[8]S. Y. Lee, M. K. Ju, H. M. Jeon, E. K. Jeong, Y. J. Lee, C. H. Kim, H. G. Park, S. I. Han, and H. S. Kang, "Regulation of tumor progression by programmed necrosis," Oxidative Medicine and Cellular Longevity **2018** (2018).

[9]H. H. Billett, "Hemoglobin and hematocrit," Clinical Methods: The History, Physical, and Laboratory Examinations. 3rd edition (1990).

[10]C. E. Rhodes, D. Denault, and M. Varacallo, "Physiology, oxygen transport," (2019).

[11]J.-A. Collins, A. Rudenski, J. Gibson, L. Howard, and R. O'Driscoll, "Relating oxygen partial pressure, saturation and content: the haemoglobin–oxygen dissociation curve," Breathe **11**, 194–201 (2015).

[12]M. O. Stefanini, F. T. Wu, F. Mac Gabhann, and A. S. Popel, "A compartment model of VEGF distribution in blood, healthy and diseased tissues," BMC Systems Biology **2**, 1–25 (2008).

[13] A. Krogh, "The rate of diffusion of gases through animal tissues, with some remarks on the coefficient of invasion," The Journal of Physiology **52**, 391–408 (1919).

[14] A. R. Anderson and M. A. Chaplain, "Continuous and discrete mathematical models of tumor-induced angiogenesis," Bulletin of Mathematical Biology **60**, 857–899 (1998).

[15] M. Elitaş and S. Zeinali, "Modeling and simulation of egf-csf-1 pathway to investigate glioma-macrophage interaction in brain tumors," International Journal of Cancer Studies & Research (IJCR): Special Issue On" Advances in Brain Cancer Research" **5**, 1–8 (2016).

[16] S. L. Chang, S. P. Cavnar, S. Takayama, G. D. Luker, and J. J. Linderman, "Cell, isoform, and environment factors shape gradients and modulate chemotaxis," PloS One **10**, e0123450 (2015).

[17] L. M. Wakefield, T. S. Winokur, R. S. Hollands, K. Christopherson, A. D. Levinson, M. B. Sporn, *et al.*, "Recombinant latent transforming growth factor beta 1 has a longer plasma half-life in rats than active transforming growth factor beta 1, and a different tissue distribution." The Journal of Clinical Investigation **86**, 1976–1984 (1990).

[18] A. Parihar, T. D. Eubank, and A. I. Doseff, "Monocytes and macrophages regulate immunity through dynamic networks of survival and cell death," Journal of Innate Immunity **2**, 204–215 (2010).

[19] M. E. Gonzalez-Mejia and A. I. Doseff, "Regulation of monocytes and macrophages cell fate," Frontiers in Bioscience **14**, 2413–31 (2009).

[20] Calculator Academy Team, "Protein hydrodynamic radius calculator,".

[21] Pubchem, "Vactosertib compound summary," (2024).

[22] S. Y. Jung, S. Hwang, J. M. Clarke, T. M. Bauer, V. L. Keedy, H. Lee, N. Park, S.-J. Kim, and J. I. Lee, "Pharmacokinetic characteristics of vactosertib, a new activin receptor-like kinase 5 inhibitor, in patients with advanced solid tumors in a first-in-human phase 1 study," Investigational New Drugs **38**, 812–820 (2020).

[23] Samsung Medical Center (Responsible Party), "Vactosertib with nal-iri/fl in metastatic pancreatic ductal adenocarcinoma," (2024).
